# A historically balanced locus under recent directional selection in responding to changed nitrogen conditions during modern maize breeding

**DOI:** 10.1101/2022.02.09.479784

**Authors:** Gen Xu, Jing Lyu, Toshihiro Obata, Sanzhen Liu, Yufeng Ge, James C. Schnable, Jinliang Yang

## Abstract

Understanding the patterns of selection during plant evolution and recent crop improvement processes is the central topic in plant breeding and genetics. As an essential macronutrient for plant growth and development, nitrogen (N) is a key factor in affecting plant adaptation and crop improvement. The widespread adoption of less expensive industrial N fixation has dramatically reshaped plant morphology by favoring compact maize plants to tolerant crowding stress. The associated genetic changes, however, have not been systematically studied. Here, we investigated maize inbred lines developed before and after the 1960s — the time point when inorganic N fertilizer started to be widely used for maize production. We identified a strong selective sweep exhibiting pronounced genomic differentiation between Old-Era (pre-1960s) and New-Era (post-1960s) inbred lines. Further study revealed population genetics statistics in the sweep exhibited patterns consistent with historical balancing selection. This balanced genomic interval is associated with a number of morphological, physiological, and metabolite traits related to vegetative N responses. A cluster of three glutamate receptor-like (GLR) genes is located within the region targeted by selection. Functional characterizations suggested differences in transcriptional activity of the GLR genes between the haplotypes carried by Old-Era and New-Era inbred lines likely play an essential role in mediating distinct N responses. The identification of both targets of selection and changes in the regulation of N responsive genes between maize lines developed in different eras sheds light on the N sensing and regulation pathways and paves the way to developing N resilient crops.

## Introduction

Through usually early hybridization events followed by selective breeding, about 150 wild plants have been domesticated into crops to meet human needs (1), including the major cereal crops of maize, rice, and wheat (2). Understanding the selection forces during these domestication and improvement processes has long been the central topic in plant genetics and breeding. Depending on the allele effects relative to fitness, the modes of selection in a diploid species include positive selection to increase the frequencies of advantageous alleles, negative selection to remove the deleterious alleles, or balancing selection to maintain both alleles (3). Unlike advantageous or deleterious alleles, alleles under balancing selection are not universally beneficial or detrimental, whose fitness changes with time, space, or population frequency (4). With the increasing availability of population-level genomic data, a number of studies have been conducted to understand the patterns of advantages or deleterious alleles (5–9). However, studies focusing on balancing selection in plants are limited (10–13), likely due to the balanced alleles being difficult to detect (14), which prevent the accurate evaluation of the roles that balanced alleles played during the crop domestication processes.

Nitrogen (N), as one of the essential macronutrients, its availability changes with time and space and, therefore, plays a critical role in plant adaptation and recent crop improvement. N is a major constituent of proteins, nucleic acids, chlorophyll, coenzymes, phytohormones, and secondary metabolites (15; 16). Plants take up inorganic N mainly in the forms of nitrate (NO_3_^−^) and ammonium (NH_4_^+^) from agricultural soils via specific assimilation and mobilization processes (17–20). For most cereal crops, such as maize or sorghum, achieving high yields in an intensive agricultural system requires a large quantity of supplemental N fertilizer. However, N utilized by most plants ranges from 30% to 50 % (21), resulting in N runoff in farm fields to form nitrous oxide (N_2_O) — a potent greenhouse gas that has 300 times the warming ability of carbon dioxide (CO_2_). In addition to the substantial adverse effects on natural ecosystems and global warming (22; 23), the poor N usage imposes the economic cost on farmers and reduces human life expectancies around the globe (24).

Maize (*Zea mays* ssp. *mays L*.) is a major crop grown around the world and consumes 17% N fertilizers worldwide (21). In the past, the breeding efforts in maize mainly focused on increasing grain yield, resulting in steady yield improvement over the last century (25). Prior to the 1960s (Old-Era) in the U.S. Corn Belt, the selection in maize breeding primarily occurred in nitrogen-limited agricultural systems. Subsequence to the green revolution in the 1960s (New-Era), inorganic nitrogen fertilizers became increasingly available due to the Haber-Bosch process (26) and maize selection and breeding has been mainly conducted in systems where nitrogen was not the limiting constraint on productivity or yield. The shift in the crucial environmental factor of N availability has changed the breeders’ preference to select hybrids with high planting density to take advantage of sufficient nitrogen fertilizers (27), resulting in a number of changes in physiological and morphological traits (28; 29). However, previous studies have not systematically examined the mode of selection act on the shifted nitrogen condition and to what extent the selection has reshaped the genomic architecture in affecting N responses.

In this study, we employed a set of Old-Era and New-Era maize inbred lines to evaluate genome wide signatures of selection to changing N conditions. We characterized N-related traits in field conditions under sufficient and N limited conditions in a two-year field trial and validated differences in phenotypic performance in controlled environment studies. Leveraging publicly available genomics dataset as well as newly generated phenomics, metabolomics, and transcriptomics datasets, our integrative analyses revealed a region which was historically under balancing selection because a direct target of positive selection during modern crop breeding. Functional characterization identified differences in transcriptional activity associated with differences in response to N. Our results shed light on the selection patterns of an N-associated locus and provide a potential target for developing N resilient crops in the future.

## Results

### Genome-wide selection scan identified a differentiated genomic region during recent breeding

We collected five in-field leaf physiological traits, including leaf nitrogen level, leaf chlorophyll content, leaf dry weight, leaf fresh weight, and leaf area from replicated field trials of the maize association panel (MAP) (30) grown under both conventional agronomic practices (high N, or HN) and under nitrogen-limited conditions (low N, or LN) in 2018 and 2019 (See **Materials and Methods**) using hyperspectral reflectance phenotyping (31) (**Table S1**). Among the 231 lines phenotyped, 37 have been previously classified as Old-Era inbred lines (i.e., lines developed before the 1960s) and 33 have been previously classified as New-Era inbred lines (i.e., lines developed post the 1960s), respectively (32) (**Table S2**). Old-Era inbred lines exhibited smaller differences than New-Era inbred lines for the leaf nitrogen level, leaf chlorophyll content, and leaf dry weight between plants grown in low N and high N conditions (**Figure 1A**, see **Figure S1** for other traits). In-field phenotypic data, especially the HN/LN ratios, suggested the New-Era lines were more responsive to take advantage of increased N availability at the vegetative stage, consistent with a previous study (33).

**Figure 1.**
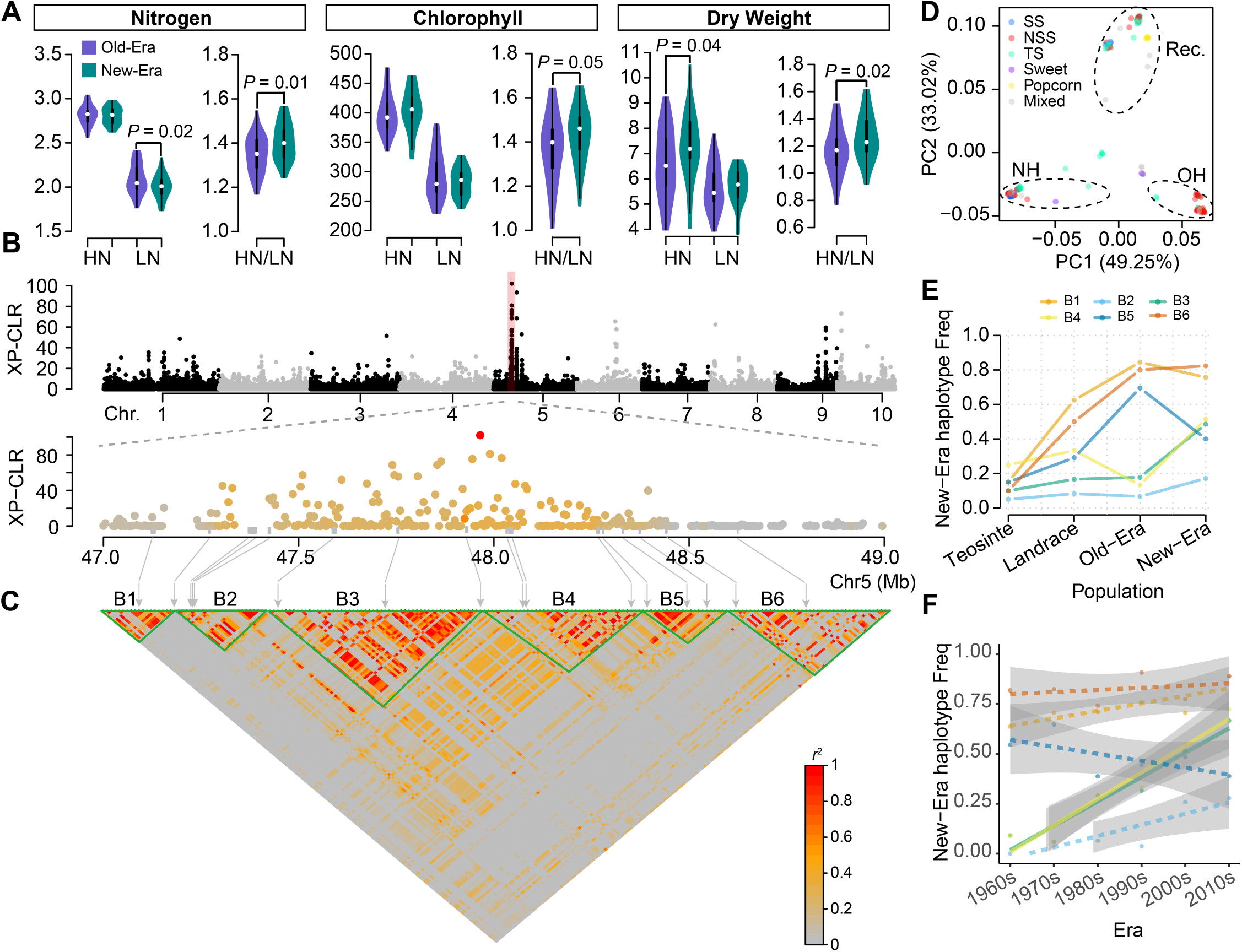
Phenomic and genomic characteristics of maize inbred lines developed before and after the 1960s. (**A**) Comparison of values for three leaf physiological traits scored across 70 maize inbreds grown under high (HN) and low N (LN) levels. (**B**) Results of a genome wide scan for selective sweeps between Old-Era and New-Era maize (see **Figure S2** for results using *F*_*ST*_ approach) including a zoomed in view of the XP-CLR scores individual windows located near the peak highlighted in red on chromosome 5. The color in the zoom-in plot reflects the LD level (*r*^2^) of windows with the leading signal (the red dot). (**C**) Linkage disequilibrium (LD) relationship for genetic markers in the highlighted region of chromosome 5. The grey arrows indicate the positions o f annotated genes. Six LD blocks (block1 to block6) are indicated with green triangles and labeled B1-B6. (**D**) PCA analysis conducted using genetic markers of 271 maize inbred lines within LD Block4. Dashed ovals indicate clusters corresponding to three haplotypes within this region, one (NH) abundant in New-Era maize, one abundant in Old-Era maize and another set of lines carrying a recombinant haplotype. Individual points are color-coded by the subpopulations of maize inbreds assigned to in Flint-Garcia et al. 2005 (30). (**E**) The frequency of the New-Era haplotype of each LD block in populations of teosinte, landrace, Old- and New-Era maize inbred lines. (**F**) Changes in the frequency of the New-Era haplotype for each LD block in elite inbred lines developed in China between the 1960s to the 2010s. Solid and dashed lines indicate significant (linear regression analysis, *P*-value < 0.001) and nonsignificant linear regressions, respectively, with 95% confidence intervals for each regression indicated in grey.

Employing both XP-CLR and *F*_*ST*_ approaches (**Materials and Methods**) to scan for signatures of selection between Old-Era and New-Era inbred lines using whole genome resequencing data resulted in the identification of 491 selective sweeps (**Table S3**). The regions identified exhibited significant overlap (75/491=15.3%, permutation test, *P*-value = 7×10^−3^) with a set of regions linked to recent sweeps associated with maize improvement (29). The most significant sweep was located on chromosome 5 and detected by both XP-CLR and *F*_*ST*_ approaches (**Figure 1B** and **Figure S2**). This sweep colocalized with genetic loci associated with nitrogen uptake efficiency (34), plant height (35), and grain weight per plant (36). We detected six linkage disequilibrium (LD) blocks (**Materials and Methods**) in the region from 1Mb upstream to 1 Mb downstream of the leading signal. These LD blocks (B2, B3, B4) exhibited pronounced genomic differentiation between New-Era and Old-Era inbred lines (**Figure 1B-C**).

For each LD block, we conducted haplotype analysis and assigned the New-Era and Old-Era haplotypes with a membership coefficient of Q *>*= 0.7 (**Materials and Methods**). As shown in **Figure 1D**, the New-Era, Old-Era, and recombinant haplotypes were each observed in maize inbreds from different subpopulations (30), suggesting population structure is unlikely to explain the observed pattern of LD (See **Figure S3** for other LD blocks). We found New-Era haplotypes of the B2, B3, and B4 LD blocks were also present at intermediate frequencies in teosinte (the maize wild ancestor) and maize landrace populations (**Figure 1E**). As expected, the frequency of the New-Era haplotypes for these three blocks were lower in Old-Era lines and then exhibit a dramatic increase in frequency after the 1960s. Consistent with the pattern observed in the MAP which composes primarily of lines developed in the Americas, in Chinese elite inbred lines (29), the frequencies of New-Era haplotypes for B3 and B4 also rose dramatically from 0.1 to 0.7 over the past 60 years (**Figure 1F**). Taken together, these data suggested the New-Era haplotype exists in the maize ancestral population and has undergone recent positive selection in both China and US elite maize populations.

#### Balancing selection maintains genetic variation at the N associated locus

In teosinte population, the B4 New-Era haplotype exhibited an intermediate frequency (> 0.2) (**Figure 1E**), higher than that in Old-Era maize inbreds, which drive us to hypothesize that this N associated locus may be under historical balancing selection (4). To address this hypothesis, we calculated the site frequency spectrum (SFS) for each LD block. Using sorghum as the ancestral alleles, we found the derived alleles, especially in the maize population, showing an excess of intermediate frequencies for B4 (**Figure 2A**), a signature consistent with balancing selection that maintains different alleles (in this case, both Old-Era and New-Era alleles) at the selected loci for a long evolutionary period (4). We observed a similar pattern for B2, B3, and B5 (**Figure S4B-D**) in the selective regions that was different from the genome-wide pattern (**Figure S4F**).

**Figure 2.**
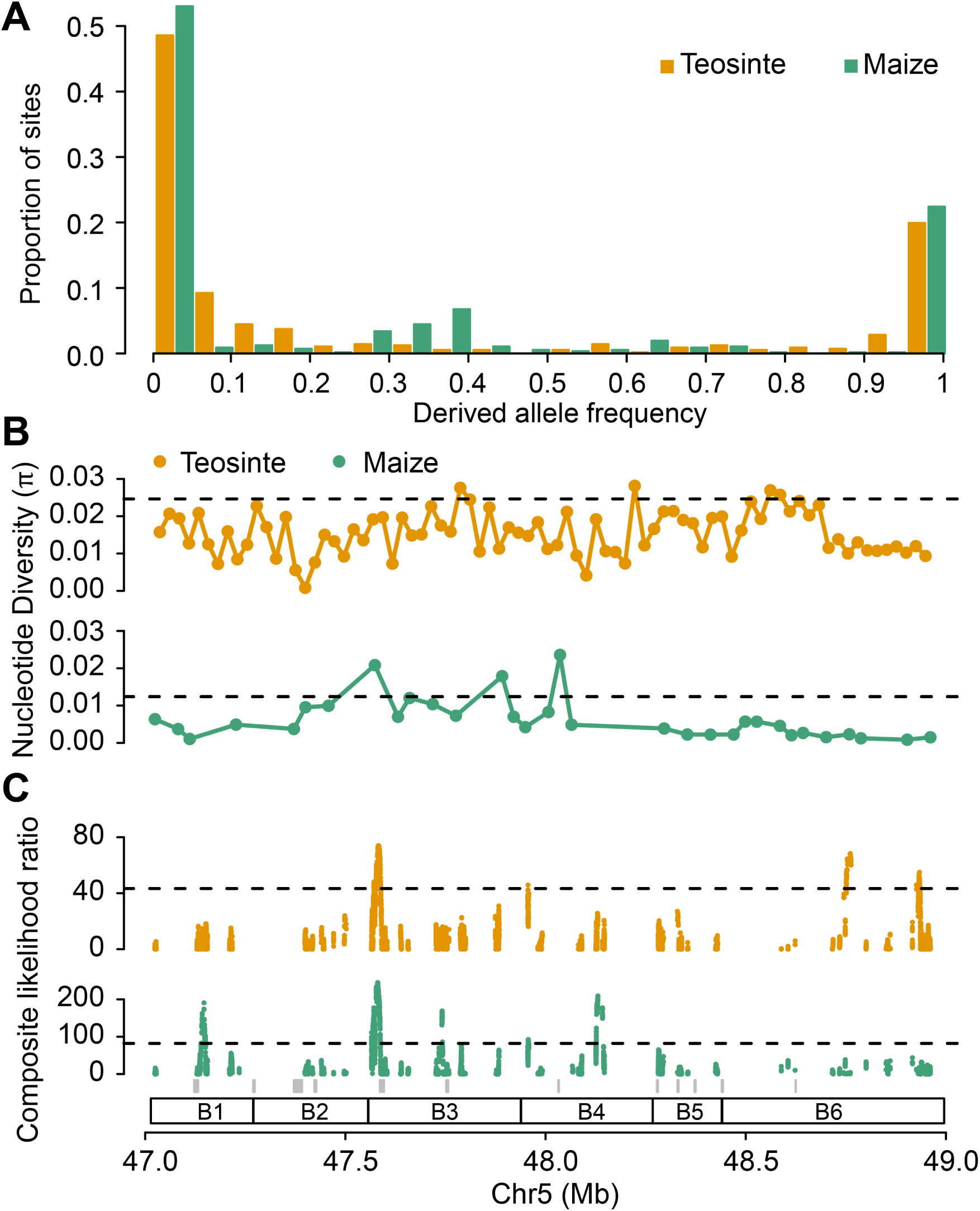
Site frequency spectrum (SFS) and neutrality test statistics at the N responsive locus. (**A**) The SFS of LD Block4 for teosinte and maize populations considering the allele shared with sorghum as the ancestral allele and the non-shared allele as the derived allele. Nucleotide diversity (*π*) (**B**) and composite likelihood ratio based on B0,*MAF* statistic (**C**) for the chromosome 5 region. The horizontal dashed lines represent the 5% significance level across the genome. The grey rectangles at the bottom of the panel C indicate the position of annotated gene models. The labels of B1 to B6 indicate the positions of the six LD blocks.

Historical balancing selection is predicted to result in high sequence diversity (4). Consistent with this model, nucleotide diversity (*π*) and Tajima’s D results (**Figure S5**) in the region are significantly higher than genome-wide level, especially within the B4 block (**Figure 2B**). Furthermore, using a newly developed composite B statistics (37), we detected balancing selection signals in the chromosome 5 region for both teosinte and maize (**Figure 2C**). These results suggest the N associated locus might be a historically balanced site to maintain for both New-Era and Old-Era alleles.

#### The selective haplotypes associated with plant morphology, physiology, and metabolite traits

Old-Era and New-Era inbreds were further characterized under controlled environment conditions in the plant phenotyping facility (**Materials and Methods**). A haplotype-based association analysis (**Materials and Methods**) detected significant differences in the N content of the lower leaves of maize lines carrying the New-Era and Old-Era haplotypes of LD B4 block (**Figure 3A**), a pattern similar to the field data under low N condition (**Figure 1A**). In contrast to the field study, leaf chlorophyll content was not significantly different under controlled environment conditions (**Figure S6**). Plants carrying the B4 New-Era haplotypes exhibited significantly larger leaf areas, greater leaf dry weights, and more compact plant architectures (**Figure 3B**). Using data on the abundance of primary metabolites collected from leaf tissue of the same plants (40), we found the abundance of lysine (C_6_H_14_N_2_O_2_), an essential amino acid, was significantly higher in inbreds carrying the B4 New-Era haplotype at the B4 linkage block (**Figure 3B**). In contrast, the abundance of fructose (C_6_H_12_O_6_) was significantly lower in inbreds carrying the New-Era haplotype at the B4 linkage block compared to inbreds carrying the Old-Era haplotype (**Figure 3B**). These differences in both leaf morphology and physiological characteristics are consistent with the view that modern maize lines were preferentially selected to take advantage of the N-oversupplied condition (41; 42).

**Figure 3.**
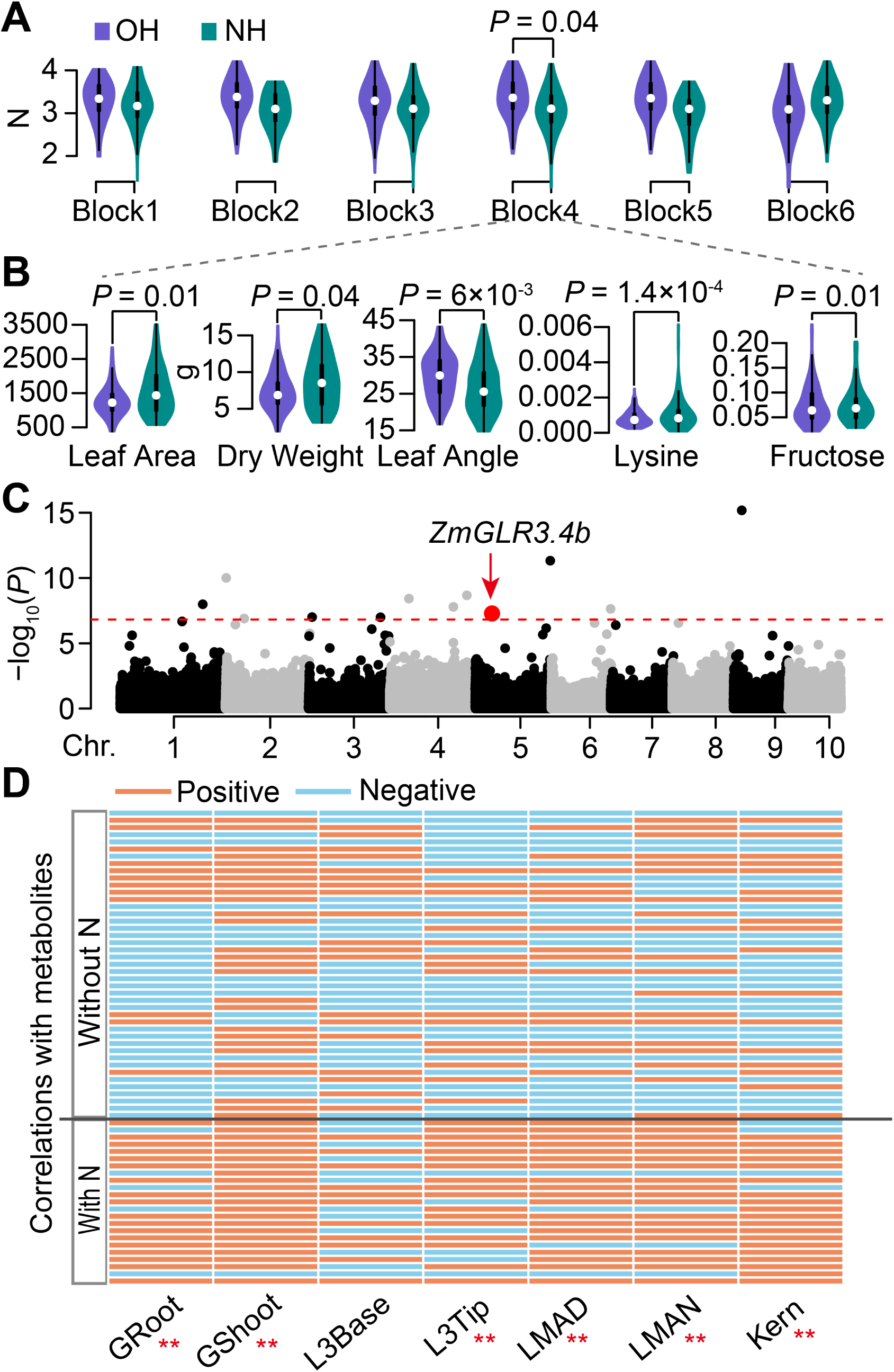
Association results for plant morphology, physiology, and metabolite traits. (**A**) Haplotype-based association analysis for leaf nitrogen level across six different LD blocks. (**B**) Phenotypic performance between Old-Era haplotype (OH) and New-Era haplotypes (OH) at LD block 4 (B4) for leaf area, leaf dry weight, leaf angle, leaf lysine content, and leaf fructose content traits. (**C**) The Manhattan plot for leaf chlorophyll *a* content using the NAM population (38). The red dot indicates the GWAS signal at chromosome 5 overlapped with *ZmGLR3.4b* gene in B4. The red horizontal dashed line denotes the Bonferroni threshold (*P* < 1.5×10^−7^). (**D**) Correlation analysis between gene expression of *ZmGLR3.4b* and 66 metabolites. “With N” and “Without N” denote the metabolites containing or not containing N in their chemical formulas. ** denotes Chi-squared test *P*-value < 0.01. Gene expression data were collected from seven tissues (39), including germinating root (GRoot), germinating shoot (GShoot), third leaf base (L3Base), third leaf tip (L3Tip), adult leaf during the day (LMAD), adult leaf during the night (LMAN), and kernel (Kern).

In total, 17 genes were annotated in the extended area of the selective sweep (**Figure 1C**). Transcriptome data revealed that 10 of these 17 genes were differentially expressed (DE) between Old-Era and New-Era inbred lines (two-sided Student’s *t*-test, *P*-value <0.05) in at least one of the seven tissues (39). These DE genes are particularly common in the LD blocks B3 (2/3) and B4 (3/3) (**Figure S7**). Among the six genes within B3 and B4, noticeably, a cluster of three glutamate receptor-like (GLR) genes was identified. GWAS for N-related traits using public data collected from the NAM population (38) identified a significant signal for variation in the abundance of chlorophyll *a* within the second exon of *Zm00001d014456*, one of the three GLR genes located within B4 (**Figure 3C**). Phylogenetic analysis of 18 maize and 20 *Arabidopsis* GLRs indicated that the cluster of three GLRs in the selective sweep was most closely related to the Arabidopsis gene *AtGLR3.4* (**Figure S8**). We refer to them below as *ZmGLR3.4a* (*Zm00001d014451*), *ZmGLR3.4b* (*Zm00001d014456*), and *ZmGLR3.4c* (*Zm00001d014458*).

The mRNA abundance of one of the three GLR genes (*ZmGLR3.4b*) exhibited a statistically significant trend towards positive correlations with the abundance of N-containing metabolites, such as lysine (C_6_H_14_N_2_O_2_), serine (C_3_H_7_NO_3_), allantoin (C_4_H_6_N_4_O_3_), gamma-aminobutyric acid (GABA, C_4_H_9_NO_2_), and negative correlations with the abundance of metabolites that do not contain the element N, such as fructose (C_6_H_12_O_6_), glyceric acid (C_3_H_6_O_4_), puruvic acid (C_3_H_4_O_3_) (**Figure 3D**). This pattern was observed when using expression data for the *ZmGLR3.4b* gene in six out of seven tissues evaluated. However, any pattern of correlation between gene expression and metabolite abundance was substantially less clear for other genes within the region (**Figure S9**).

#### Expression of ZmGLR3.4b is affected by altered cis-regulatory modulation

The overall expression levels of *ZmGLR3.4a* was much lower than that of *ZmGLR3.4b* or *ZmGLR3.4c* (**Figure S10**). Both *ZmGLR3.4b* and *ZmGLR3.4c* were predominantly expressed in leaf tissues. In the leaf three tip (L3Tip) and adult leaf collected during the day (LMAD), the expression levels of *ZmGLR3.4b* was significantly higher in New-Era than in Old-Era inbreds. In contrast, the expression of *ZmGLR3.4c* in L3Tip was significantly lower in New-Era inbred lines than in Old-Era inbreds (**Figure S10**). Genome-wide analysis identified significant *cis*-eQTL for both *ZmGLR3.4b* (**Figure 4A**) and *ZmGLR3.4c* (**Figure S11**).

**Figure 4.**
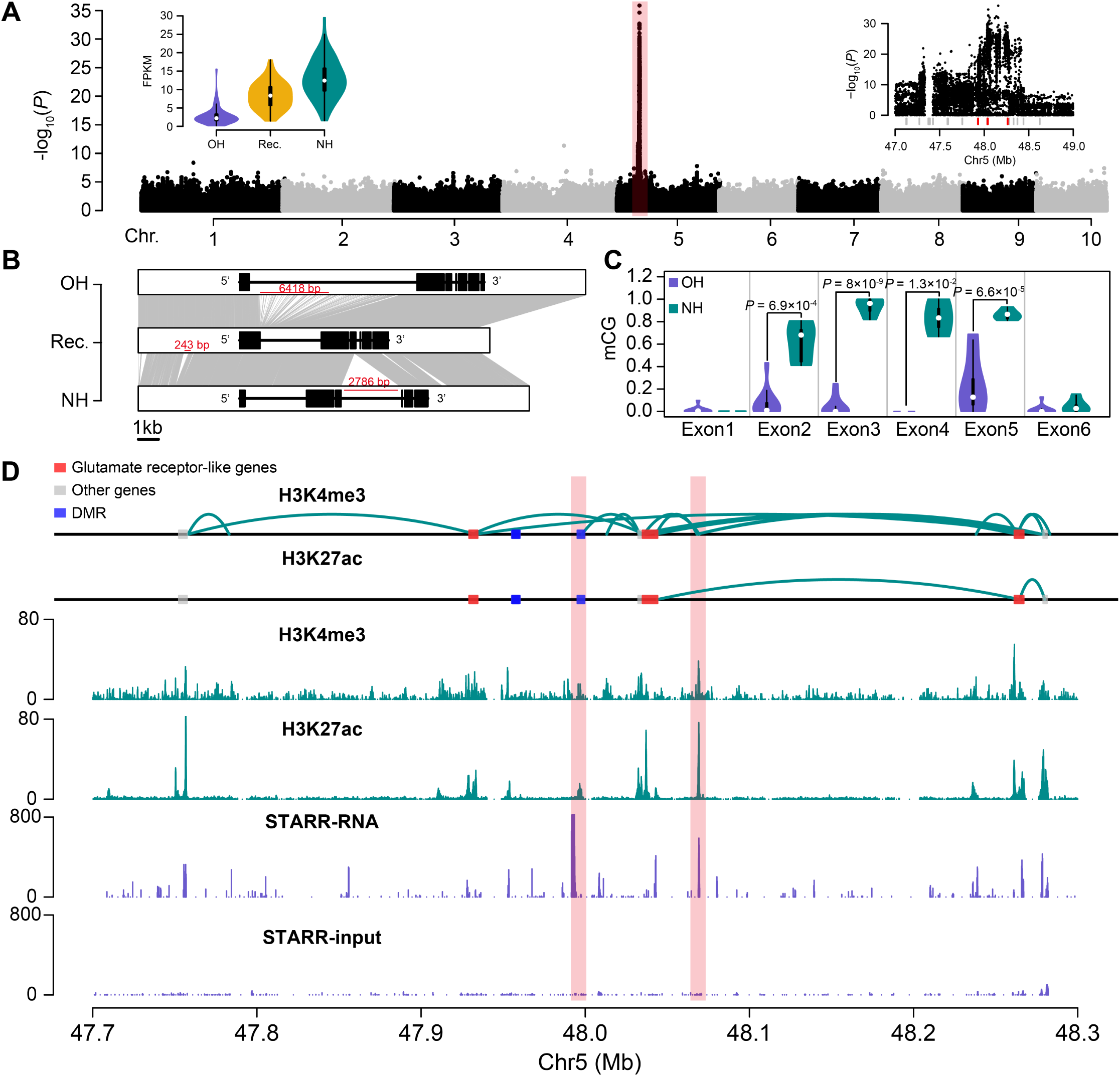
Functional genomic characterization of the GLR genes at the chromosome 5 genomic interval. (**A**) Results of a genome-wide eQTL analysis using the expression of the *ZmGLR3.4b* gene collected from third leaf tip as the trait. The distribution of the expression levels of the *ZmGLR3.4b* gene within the Old-Era (OH), New-Era (NH), and recombinant (Rec.) haplotypes are shown in the top left panel. The top right panel shows a zoom-in view of the region of the genome wide Manhattan plot highlighted in red. The positions of the three GLR genes in the top right panel are indicated by the three red tick marks. (**B**) Comparison of the annotated structure of the *ZmGLR3.4b* gene in *de novo* assembled genomes of maize lines carrying the Old-Era (OH, A188), New-Era (NH, B73), and recombinant (Rec., IL14H) haplotypes. (**C**) Levels of DNA methylation in exons of *ZmGLR3.4b* belonging to Old-Era (OH, *n* = 9) and New-Era haplotypes (NH, *n* = 5). *P* values were determined using a two-sided Student’s *t*-test. (**D**) Physical interactions (two upper panels), colocalization with H3K27ac and H3K4me3 (two middle panels), and STARR profiles (two lower panels) around the GLR genes. Curved green lines denote interacting regions with evidence of physical contact. Red and gray boxes indicate the GLR and other gene models, respectively. Blue boxes indicate teosinte-maize differentially methylated regions. The regions highlighted in pink indicate anchors showing enhancer activities.

Leveraging the *de novo* assembled maize genomes that include both Old-Era and New-Era haplotypes (43–45), we investigated the structural variation (SV) of the GLR genes. No apparent structural variation was present in the *ZmGLR3.4a* and *ZmGLR3.4c* genes between *de novo* assembled genomes for inbreds carrying the Old-Era and New-Era haplotypes (43–45) (**Figure S12**). However, two transposable element (TE) insertions, present in the first and third introns, distinguished *ZmGLR3.4b* in the genome of inbreds carrying the Old-Era and New-Era haplotypes (**Figure 4B**). The 2,786-bp TE insertion in the third intron in the New-Era haplotype, is likely associated with the spread of both CG (**Figure 4C**) and CHG (**Figure S13**) DNA methylation into the surrounding exons of the *ZmGLR3.4b* gene. Published H3K4me3 and H3K27ac ChIP-seq data (46; 47), as well as the STARR-seq (47) data, suggested that multiple putative promoters or enhancers exist at the selective region (i.e., B3 and B4). HiChIP data further illustrated physical contacts among the three GLR genes in B73, an inbred line carrying the New-Era haplotype (**Figure 4D**). In addition, *ZmGLR3.4b* physically interacted with two putative promoters or enhancers (highlighted areas in **Figure 4D**), as evidenced by the high STARR-seq or ChIP-seq peaks; one of the physical interactions overlaps with a differentially methylated region (DMR) that was previously identified between maize and teosinte (48). Taken together, these results suggest *cis*-regulatory modulations likely alter the transcriptional activities of the GLR genes.

#### Phenotypic and transcriptional responses under different N conditions

We conducted additional growth chamber experiments using four selected inbred lines (two lines for each Era) based on their haplotypes and field performance. After growing for two weeks with different N treatments, we harvested the aboveground and belowground tissues for phenotyping (**Materials and Methods**). The dry weights of both aboveground and belowground tissue of Old-Era lines were not significantly different between plants grown in high N and low N treatments (**Figure 5A**). A similar phenomenon was observed in the fresh weight (**Figure S14**), consistent with the N resilient effect of the Old-Era haplotype observed in the field. For the New-Era lines, both genotypes exhibited significantly better performances under high N as compared to low N conditions, except for the dry root weight of B73, consistent with previous observations that the New-Era haplotypes were more responsive to N.

**Figure 5.**
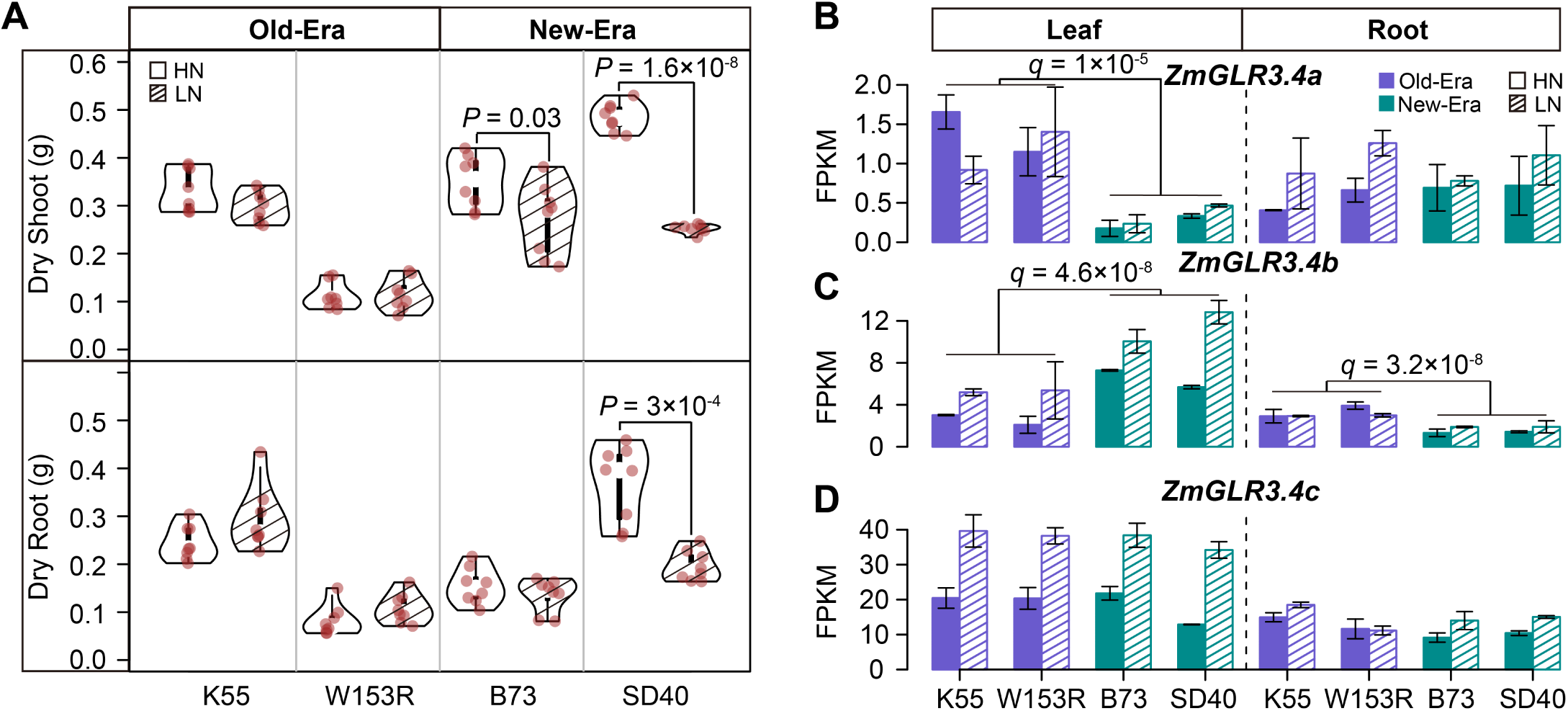
Phenotypic and transcriptional responses of selected Old-Era and New-Era inbred lines to different N treatments. (**A**) The dry weight of two-week-old shoot and root for Old-Era and New-Era inbred lines growing with high N (HN) and low N (LN) treatments. *P* values were determined using a two-sided Student’s *t*-test. Gene expression of *ZmGLR3.4a* (**B**), *ZmGLR3.4b* (**C**), and *ZmGLR3.4c* (**D**) of Old-Era and New-Era inbred lines with different N treatments. FDR corrected *P*-value (*q*) was calculated between Old-Era and New-Era inbred lines.

The expression of *ZmGLR3.4a* was consistently low (FPKM *<* 2) in both public and newly generated RNA-seq data from the plants grown in this study (**Table S4**), indicating it is a potentially malfunctional gene (**Figure 5B**). In leaf tissue, the abundance of mRNA transcripts derived from both *ZmGLR3.4b* (**Figure 5C**) and *ZmGLR3.4c* (**Figure 5D**) responded positively to high N treatments; in root, no apparent transcriptional reactions to the N treatments were observed for either gene. New-Era lines exhibited significantly higher expression of *ZmGLR3.4b* in leaf tissue than did Old-Era lines (fold change = 2.2, FDR corrected *P*-value or *q*-value = 4.6×10^−8^) under both high N (fold change = 2.5, *q*-value = 1.5×10^−8^) and low N (fold change = 2.1, *q*-value = 1.7×10^−3^) conditions (**Figure 5C**). In root tissue, the opposite pattern was observed with a moderate decrease in the expression of *ZmGLR3.4b* New-Era inbred lines relative to Old-Era inbred lines (fold change = 2.4 for HN and fold change = 1.5 for LN). We did not detect any significant differences in the expression of *ZmGLR3.4c* between Old-Era and New-Era inbred lines (**Figure 5D**).

We detected *n* = 2,264 differentially expressed (DE) genes in leave tissue and *n* = 699 DE genes in root tissue, comparing expression between the two N treatments (**Table S5**). These genes are referred to below as N responsive genes. Additionally, we detected *n* =1,600 leaf and *n* = 1,609 root DE genes between Old-Era and New-Era inbreds (**Table S5**). Notably, the Old-Era vs. New-Era DE genes were significantly enriched in the N responsive gene sets (**Figure S15**). KEGG analysis suggested the DE genes were highly enriched in metabolism pathways (**Figure S16**). Old-Era vs. New-Era DE genes were enriched in genes encoding enzymes from multiple amino acids metabolism pathways, including the glutamate metabolism pathway (**Figure S16**). Plant GLRs are largely glutamate non-specific and can be gated by a broad range of amino acids (49). To gain further insight into the roles of the *ZmGLR3.4* genes, we employed the predicted protein-protein interaction (PPI) networks using New-Era and Old-Era DE genes in the leaf tissue. After pulling down the network involving *ZmGLR3.4a* and *ZmGLR3.4b* (*ZmGLR3.4c* was not differentially expressed between Old-Era and New-Era inbreds as shown in **Figure 5D**), we found *ZmGLR3.4* genes positioned in between the N synthesis and transportation pathways and the ion signaling pathways (**Figure 6**). Noticeably, a number of known N assimilation and transportation genes, i.e., *ZmNIR1.1* (*Zm00001d052165*) (50), *ZmNR1.1* (*Zm00001d018206*) (50), *ZmAMT1.1B* (*Zm00001d003025*) (51), were up-regulated in New-Era lines, while the ion transporters, i.e., *ZmHKT1* (*Zm00001d040627*) (52), *cbl4* (*Zm00001d038730*), *ZmHAK25* (*Zm00001d017925*) (53), were down-regulated.

**Figure 6.**
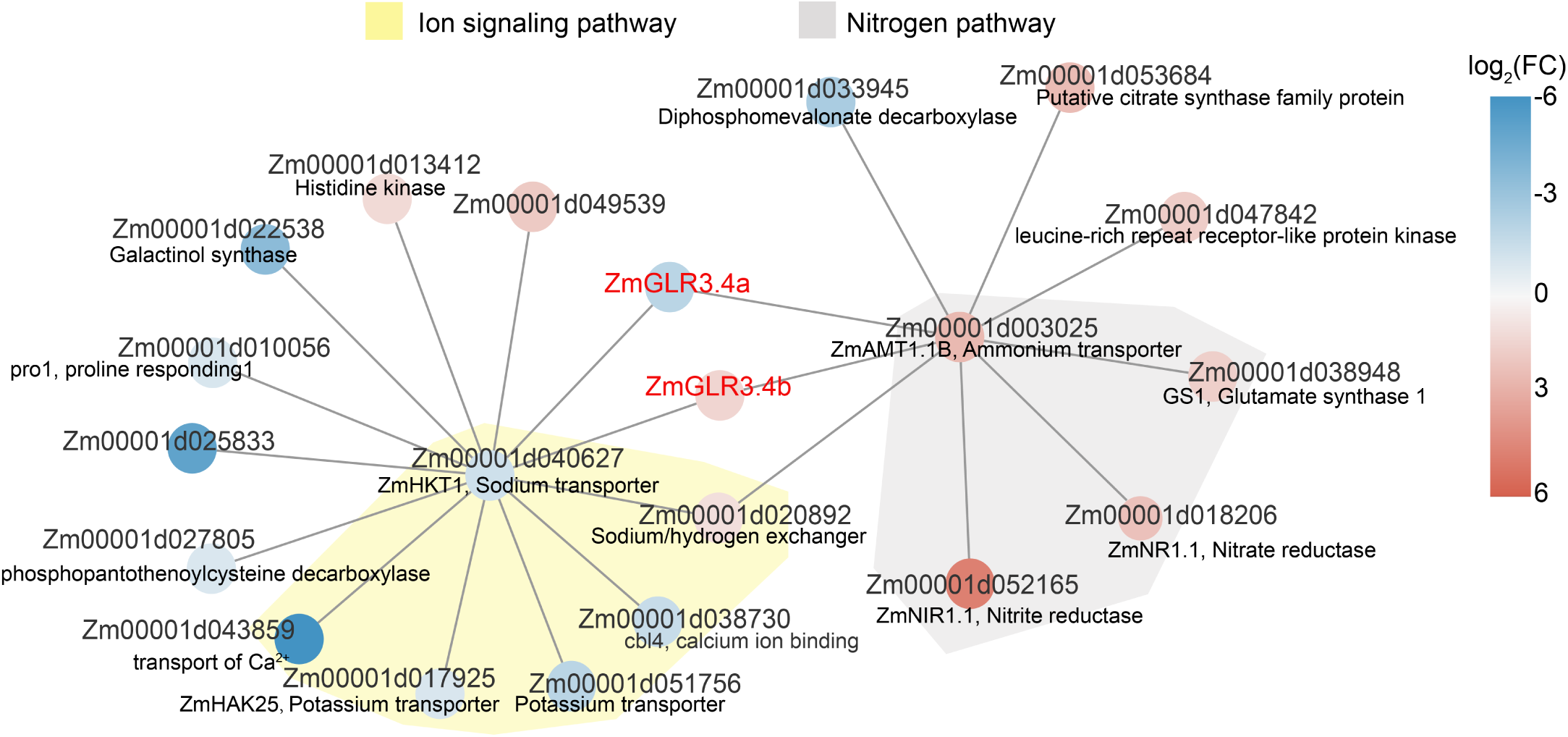
Network analysis of differentially expressed (DE) genes between Old-Era and New-Era inbreds. Protein-protein interaction network predicted from Old-Era vs. New-Era DE genes detected in the leaf tissue. The gradient colors of the dots denote the log2 fold change of NE/OE ratio. The yellow and gray colors shaded areas highlight genes involved in ion signaling and nitrogen pathways, respectively.

## Discussion

In this study, we found an N-associated locus that remained at an intermediate frequency from teosinte to landrace but rapidly increased its frequency in maize inbred lines developed after the 1960s when inorganic N fertilizer became increasingly available in maize production. This genetic locus is associated with vegetative N-related traits, as evidenced by field, greenhouse, and growth chamber experiments. Noticeably, the New-Era haplotype is more responsive to high N conditions, and the Old-Era haplotype is more resilient to N stress. Population genomics analyses detected an excess of SNPs with intermediate frequencies, and the sweep showed slightly increased nucleotide diversity. Additionally, a weak balancing selection signal was detected in the region. Because we only considered SNPs for the balancing selection scan, the actual sequence diversity might be underestimated due to large insertion or deletion polymorphisms. By leveraging the recently *de novo* assembled maize genomes that include both Old-Era and New-Era haplotypes, structural variation (SV) analyses found two transposon insertions in the first and third introns of the *ZmGLR4.3b* gene. Both transposable elements are historical insertion events as their sequences have been largely degraded, consistent with the idea that long-term balancing selection might maintain both Old-Era and New-Era haplotypes.

Notably, three glutamate receptor-like (GLR) genes in a tandem array were located within the selected region. The GLR genes, plant homologs of mammalian ionotropic glutamate receptors (iGluR), have been hypothesized to play a crucial role in sensing the amino acid level at the cellular level (17). Our data revealed that the gene expression levels of *ZmGLR4.3b*, the strongest candidate in the GLR gene cluster, are positively correlated with the abundance of primary metabolites containing N and negatively correlated with metabolites lacking N element. These correlations are consistent with the potential role of the GLR genes in regulating the N/C metabolic balance, as suggested by studies in Arabidopsis (54; 55). Unlike in mammals, the plant GLR genes were reported as broad-spectrum amino acid receptors (56; 57). Glutamine and glutamate, as the products of the N assimilation, are precursors of other amino acids (58; 59). The binding of GLR with amino acid likely induces a conformational change and opens the ion channel (60). Consistent with this idea, our PPI network analysis positioned GLR genes between two groups of functional genes, one group for nitrogen biosynthesis or transportation and the other for ion signaling exchanging or transportation. After ions pass through the channel, the ion signaling can potentially feedback in regulating N uptake and utilization (17; 49; 57). Note that the known N uptake and transportation genes in the PPI network were upregulated in New-Era inbred lines, likely in responding to the ion signaling feedback.

Collectively, our results suggest a functional role of the GLR genes in N responses. Population genomics analyses show signs of historical balancing selection for a selective sweep region harboring a cluster of three GLR homologous genes. In addition, the GWAS study detects genomic variation associated with several N-related traits. Around the GLR genes, we also identify functional variations, i.e., SV and DMRs, and other genomic features important for gene activation or chromatin interaction. These results suggest that further investigation of the GLRs in affecting N-related traits is warranted and that Old-Era alleles may provide a promising alternative allele for tolerating N stress and developing N resilient crops.

## Materials and Methods

### Field experimental design

The Maize Association Panel (30) including Old-Era and New-Era inbred lines were grown at the Havelock Research Farm of the University of Nebraska-Lincoln using an incomplete split plot block design in 2018 and 2019. Two nitrogen treatments were applied to the association panel, one under low nitrogen condition (no additional N fertilizer) and the other following conventional N application practice at the rate of 135 kg/ha dry urea. Each treatment was replicated twice in each year, with each replicate consisting of 288 plots including both the lines of the Maize Association panel and between 27 and 37 plots planted with check varieties. Each plot was 1.6 m wide and 6.3 m long, comprising of two rows, 38 plants were grown in each row. In-field vegetative N related traits were quantified follow a high-throughput phenotyping protocol described previously (31).

### Calculation of the best linear unbiased predictions for the field phenotypic data

To minimize the effects of environmental variation, best linear unbiased predictions (BLUPs) were performed using the R package lme4 (61) to estimate the phenotypic value. The BLUP model was:

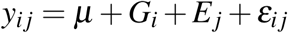

where *y*_*i j*_ is the observed phenotype for the *i*^*th*^ genotype of the *j*^*th*^ environment, *µ* is the overall mean, *G*_*i*_ is the random genotypic effect of the ith genotype, *E*_*j*_ is the random effect of the jth environment, and *ε*_*i j*_ is a random residual error.

### Plant materials and growth conditions in the controlled environments

We conducted nitrogen treatment experiments in the growth chamber with a photoperiod of 16/8 h at 28/24°C (light/dark). Seeds of two Old-Era inbreds (K55 and W153R) and two New-Era inbreds (SD40 and B73) were sterilized in 75% (v/v) ethanol for 15 min, washed with distilled water, and then soaked in distilled water for 8 h. Afterward, two seeds were planted in a plastic pot consisting of an autoclaved mixture (volume based) of 50% medium size (0.5–0.3 mm) commercial grade sand, 30% fine vermiculite, 20% MetroMix 200. Two days before planting, each tray containing 12 pots was irrigated with 3 L of a nutrient solution adjusted to pH 5.8. The high-N treatment nutrient solution contained 6.5 mM KNO_3_, 4 mM Ca (NO_3_)_2_·4H_2_O, 1 mM NH_4_H_2_PO_4_, 2 mM MgSO_4_·7H_2_O, 46 M H_3_BO_3_, 9 M MnCl_2_·4H_2_O, 7 M ZnSO_4_·7H_2_O, 0.8 M Na_2_MoO_4_·2H_2_O, 0.32 M CuSO_4_·5H_2_O, 77 M Fe-EDTA. The low-N treatment nutrient solution contained 0.65 mM KNO_3_, 4.95 mM KCl, 0.40 mM Ca (NO_3_)_2_·4H_2_O, 3.60 mM CaCl_2_·2H_2_O, 0.10 mM NH_4_H_2_PO_4_, 0.90 mM KH_2_PO_4_, 2 mM MgSO_4_·7H_2_O, 46 M H_3_BO_3_, 9 M MnCl_2_·4H_2_O, 7 M ZnSO_4_·7H_2_O, 0.8 M Na_2_MoO_4_·2H_2_O, 0.32 M CuSO_4_·5H_2_O, 77 M Fe-EDTA. On day 5, each pot was thinned to one plant and received 50 mL of high-N or low-N nutrient solution every two days. After two weeks, the shoots and roots were harvested for further experiments.

### Sequence Alignment and Phylogenetic Analysis

We downloaded the GLR protein sequences of *Arabidopsis thaliana* from the TAIR database (http://www.arabidopsis.org/), which have been reported in a previous study (62). Then the *Arabidopsis* GLR family protein sequences were aligned to the maize protein database using BLASTP (63) with an e-value of 10^−5^ to get a list of hits against maize protein. The GLR protein sequences of *Arabidopsis* as well as their orthologous genes in maize were aligned with MUSCLE (64) using MEGA6 software (65). A neighbor-joining (NJ) method was then used for phylogenetic tree construction, with 1,000 bootstrap resampling.

### Linkage disequilibrium (LD) analysis and haplotype construction

We estimated LD with the *r*^2^ statistics using plink1.9 (66). Heat maps of pairwise LD between SNPs were plotted using the R package LDheatmap (67). The R package gpart (68) was used to partition the sweep region into blocks. Haplotypes of each block was determined by Admixture (69), individuals with membership coefficients of Q *>*= 0.7 were assigned to a specific haplotype. We defined the haplotypes from New-Era and Old-Era inbreds as New-Era and Old-Era haplotypes, respectively. Haplotype based target association mapping was performed for each block using the first three principal components as the covariates in the model.

### Genome-wide association study

To determine the contribution of regulatory variants that influence gene expression of *ZmGLR3.4* genes, we conducted eQTL analysis using a mixed linear model (MLM) implemented in GEMMA (v 0.98.3) (70). The genotype was downloaded from maize HapMap 3 (71) and gene expression data was obtained from Kremling et al., 2018. The kinship matrices and the first three principal components were calculated by GEMMA (70) and Plink 1.9 (66), respectively and then fitted into the GWAS model. GWAS for leaf chlorophyll *a* content in the NAM population (38) was performed using FarmCPU method implemented in the R package rMVP (72). The NAM genotypic data was downloaded from the Panzea website (http://www.panzea.org).

### Genome-wide scanning to detect selective signals

We performed genome scanning for selective signals using *F*_*ST*_ (73) and the latest version of XP-CLR (74) approaches. In the XP-CLR analysis, we used a 50 kb sliding window and a 5 kb step size. To ensure comparability of the composite likelihood score in each window, we fixed the number assayed in each window to 200 SNPs. In the *F*_*ST*_ analysis, we calculated *F*_*ST*_ using VCFtools (75) with a 50 kb sliding window and a 5 kb step size. We merged nearby windows falling into the 10% tails into the same window. After window merging, we considered the 0.1% outliers as the selective sweeps.

### Detection of balancing selection

According to the previous study (76; 77), the locus under balancing selection has an elevated nucleotide diversity (*π*), Tajima’s D, and site frequency spectrum. Utilizing Sorghum bicolor alleles as the ancestral state, we calculated the site frequency spectrum for each block at the selection region. The nucleotide diversity (*π*) and Tajima’s D were estimated using ANGSD software (78). All bam files of 17 teosinte, 23 landrace and 269 modern maize lines were derived from the Maize HapMap3 panel (71) which were downloaded from CyVerse database (/iplant/ home/shared/panzea/raw_seq_282/bam/) as described in Panzea database (www.panzea.org). In the analysis, we first inferred the unfolded genome-wide site frequency spectrum (SFS) and calculated the thetas for each site. We then used the “thetaStat” program, which was implemented in ANGSD, to summarize the nucleotide diversity and Tajima’s D values with 25 kb non-overlapping sliding windows. We also calculated the B_0,*MAF*_ statistics using BalLeRMix (version 2.3) (37) with default parameters.

### RNA sequencing and data analysis

Two Old-Era inbreds (K55 and W153R) and two New-Era inbreds (SD40 and B73) that were planted in the growth chamber were used for RNA-seq. For each genotype, we conducted high nitrogen and low nitrogen treatments. For each treatment, we used two biological replicates for conducting RNA-seq. Two weeks after sowing, shoot and root were harvested. Total RNA was extracted and purified using the RNeasy Plant mini kit (Qiagen), followed by DNase treatment. We sequenced the libraries using Illumina NovaSeq 6000 with 150 bp paired-end reads. Raw reads were trimmed and preprocessed using fastp (79) in default settings. Using the “Rsubread” software package (80), all clean reads were then mapped to the B73 reference sequence (AGPV4) (81; 82) by the “align” function, and transcript counts were extracted by using the “featureCounts” function. We carried out differential gene expression analysis using DESeq2 (83). A gene was identified as differentially expressed if the false discovery rate (FDR) is < 0.05 and have at least twofold expression change. The expression of each gene was normalized to fragments per kilobase of transcript per million reads (FPKM).

### Pathway enrichment and protein-protein interaction analysis

We performed pathway enrichment analysis using KOBAS v2.0 (84) with a significance cutoff of *P*-value < 0.01. Protein–protein interaction (PPI) networks was established using STRING v11 with a combined score >= 0.4 (85) and visualized using Cytoscape app (86).

## Data and code availability

RNA-seq data produced from this study have been deposited in the NCBI GEO database under PRJNA800008. The code used for the analyses can be accessed through GitHub repository (https://github.com/GenXu1/Nitrogen_project).

## Acknowledgements

We thank J. Ross-Ibarra for comments on the manuscript. This work is supported by the Agriculture and Food Research Initiative Grant number 2019-67013-29167 from the USDA National Institute of Food and Agriculture, the National Science Foundation under award number OIA-1557417 for Center for Root and Rhizobiome Innovation (CRRI), and the University of Nebraska-Lincoln Start-up fund. This work was conducted using the Holland Computing Center of the University of Nebraska-Lincoln, which receives supports from the Nebraska Research Initiative.

## Author contributions

J.Y. designed this work. J.L., T.O., Y.G., J.C.S. generated the data. G.X., J.L., and J.Y. analyzed the data. S.L. provided conceptual advice. J.Y. and G.X. wrote the manuscript.

## Supporting Information

### Supporting Tables

**Table S1**. The best linear unbiased predictors (BLUP) values of leaf nutrients in maize association panel. (https://github.com/GenXu1/Nitrogen_project/tree/main/table/TableS1_field_phenotype.xlsx)

**Table S2**. Old- and New-Era samples. (https://github.com/GenXu1/Nitrogen_project/tree/main/table/TableS2_Old_New_samples.xlsx)

**Table S3**. Selective sweeps detected between Old- and New-Era inbreds. (https://github.com/GenXu1/Nitrogen_project/tree/main/table/TableS3_Sweeps.xlsx)

**Table S4**. Summary for RNA-Seq data. (https://github.com/GenXu1/Nitrogen_project/tree/main/table/TableS4_RNA_seq_data_summary.xlsx)

**Table S5**. Differentially expressed (DE) genes detected in this study. (https://github.com/GenXu1/Nitrogen_project/tree/main/table/TableS5_differentially_expressed_genes.xlsx)

### Supporting Figures

**Figure S1.**
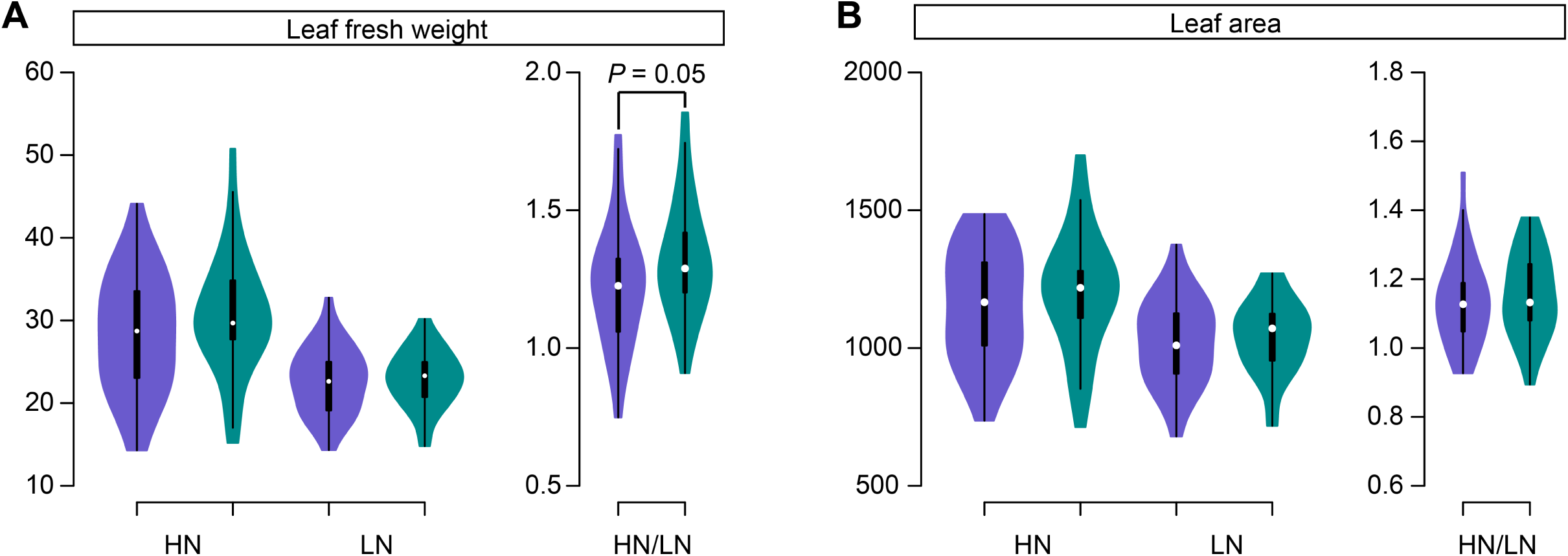
In-field phenotypic performance of Old-Era and New-Era inbred lines. Leaf fresh weight (**(A)**) and leaf area (**(B)**) under high N and low N conditions, and the ratio of high N over low N.

**Figure S2.**
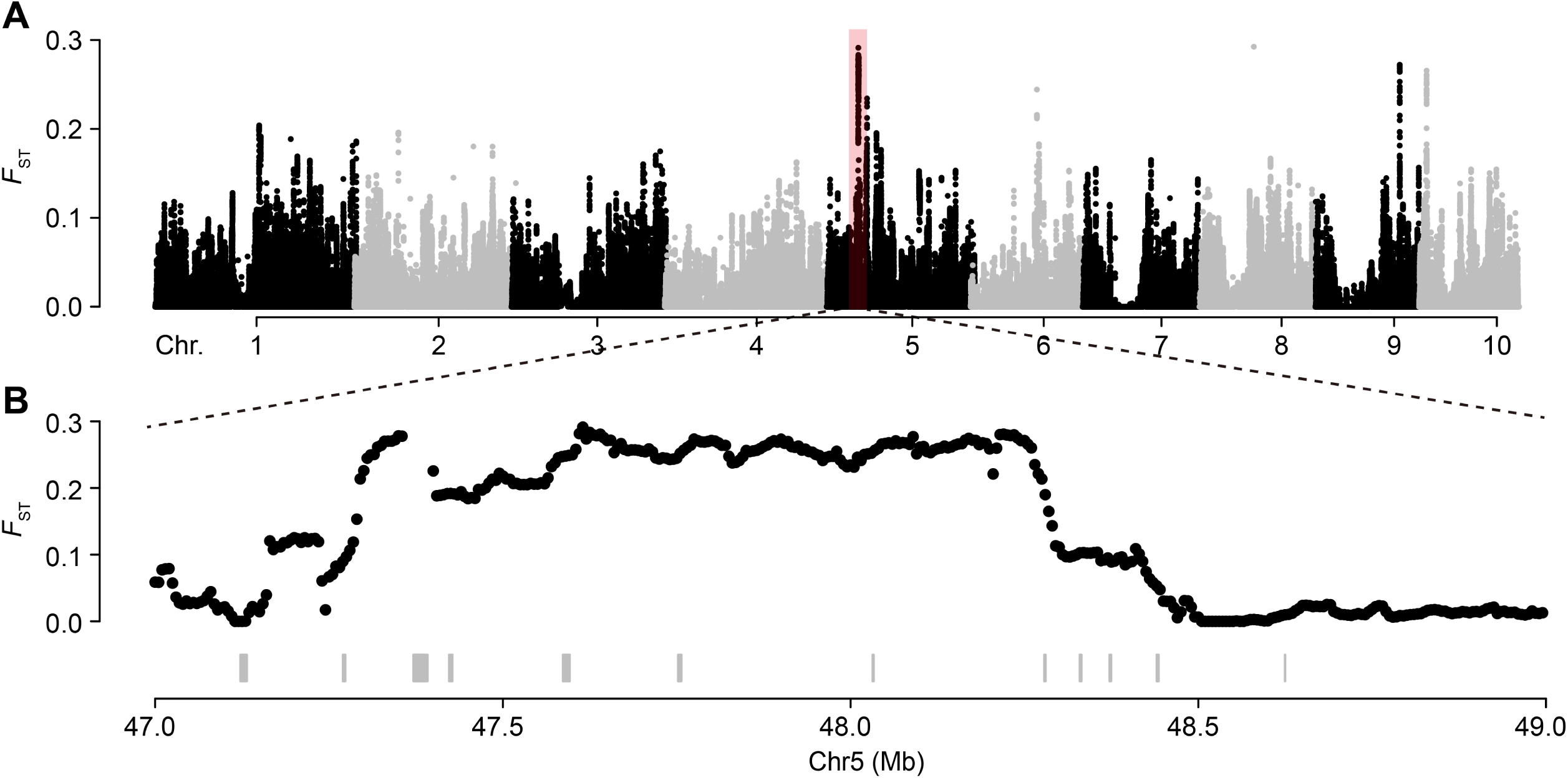
Selective sweeps detected using *F*_*ST*_ approach. Genome-wide *F*_*ST*_ values **A** and the zoom-in plot of the highlighted region on chromosome 5 **B**.

**Figure S3.**
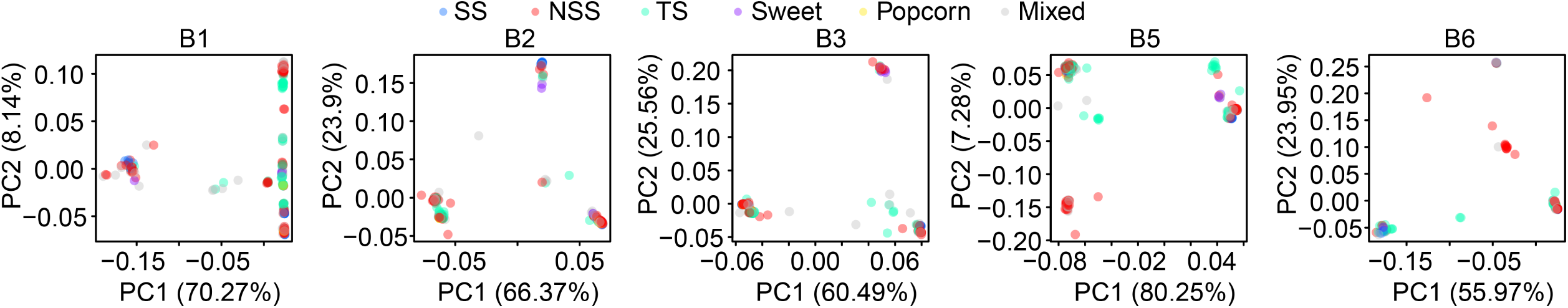
The principal component analysis (PCA) plots for linkage disequilibrium (LD) blocks. Colors represent subpopulations based on Flint-Garcia et al. 2005. B1, B2, B3, B5, and B6 denotes LD block1, block2, block3, block5 and block6, respectively.

**Figure S4.**
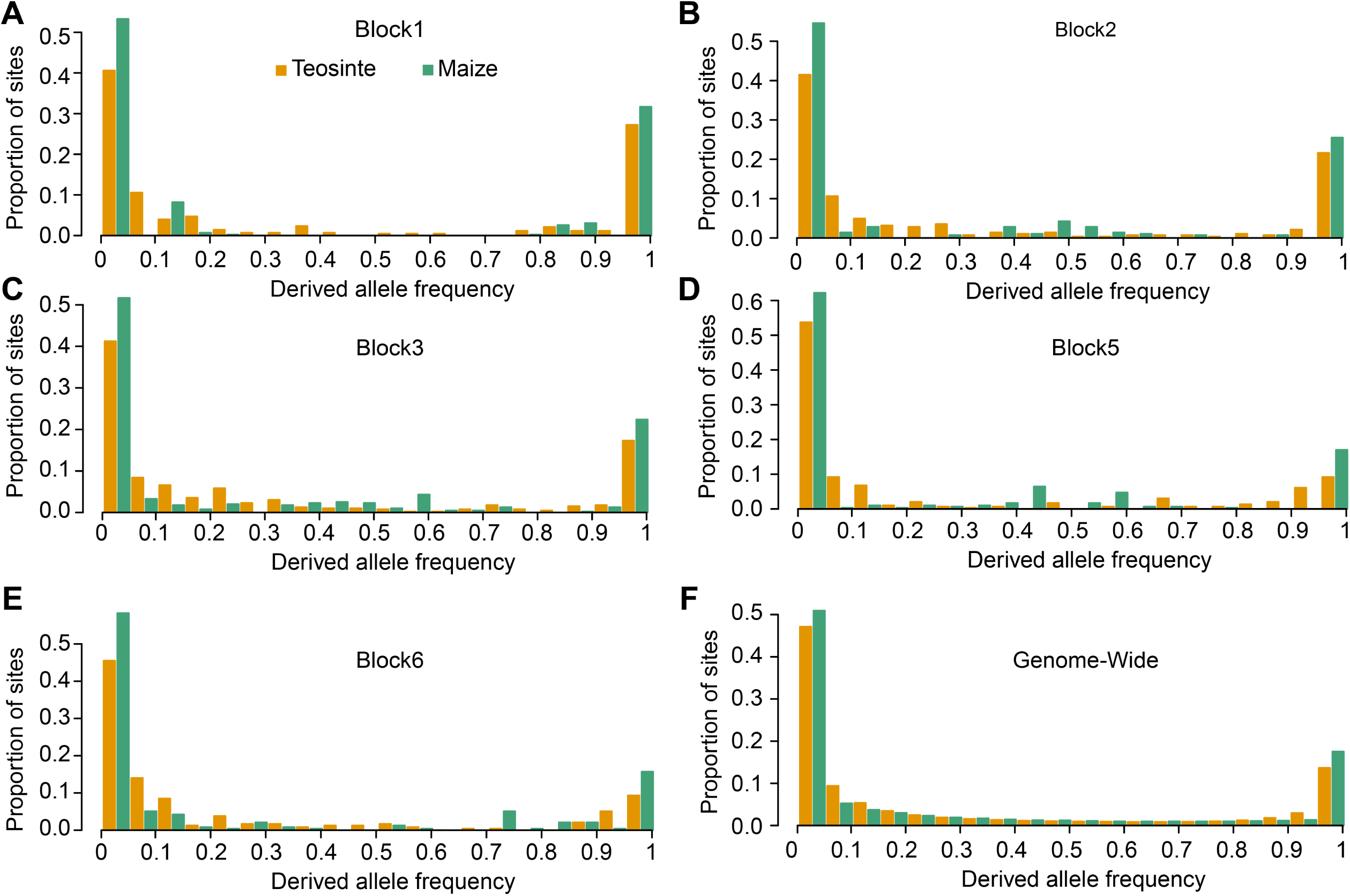
Unfolded site frequency spectrum in teosinte and maize. **(A-E)** The site frequency spectrum of variants within the LD Blocks. **(F)** Genome-wide site frequency spectrum. Sorghum bicolor alleles were used as the ancestral state.

**Figure S5.**
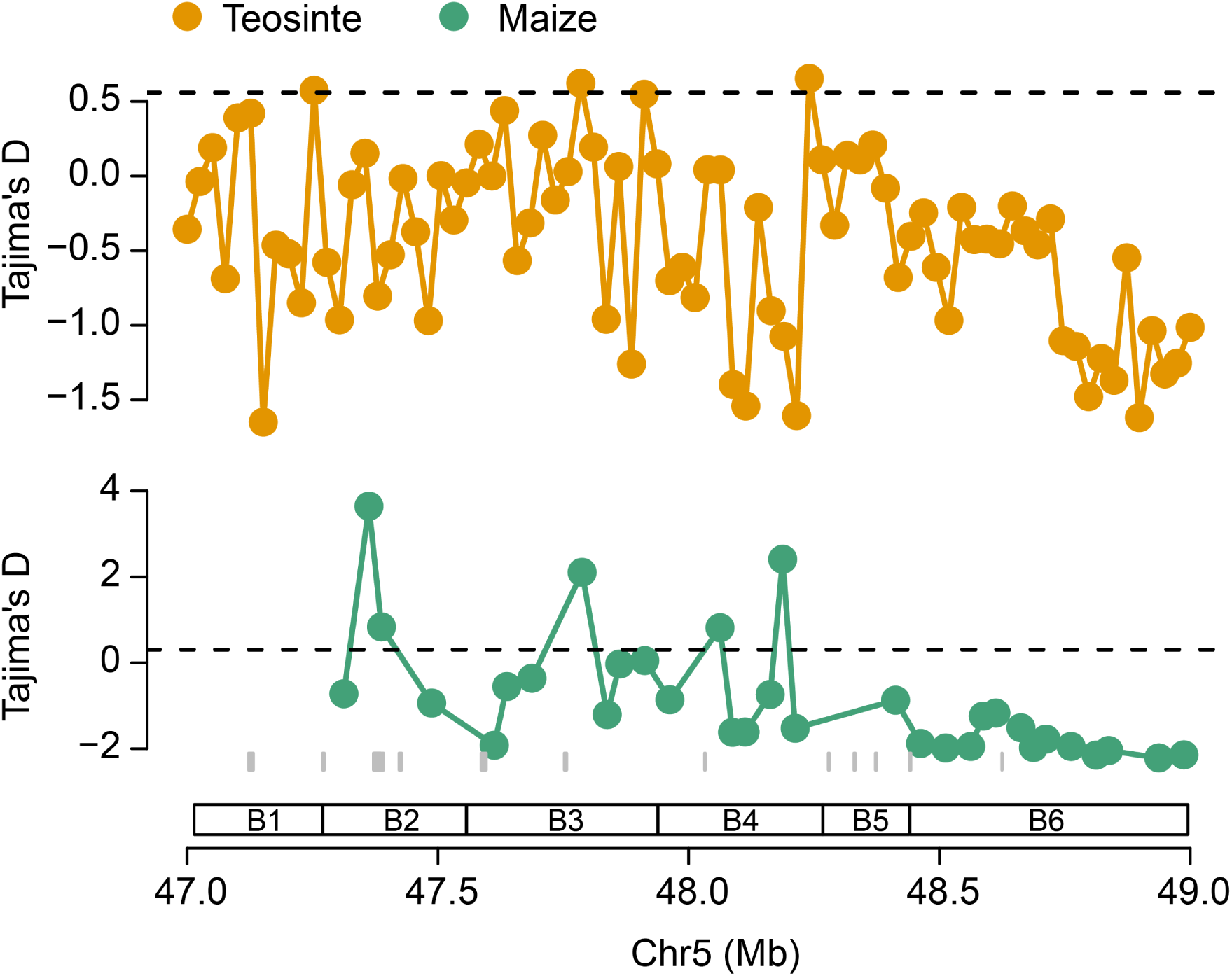
Tajima’s D at the N responsive locus on chromosome 5. The horizontal dashed lines represent the 5% significance level across the genome. The underneath grey rectangles represent gene models. B1 to B6 indicate LD Block1 to Block6.

**Figure S6.**
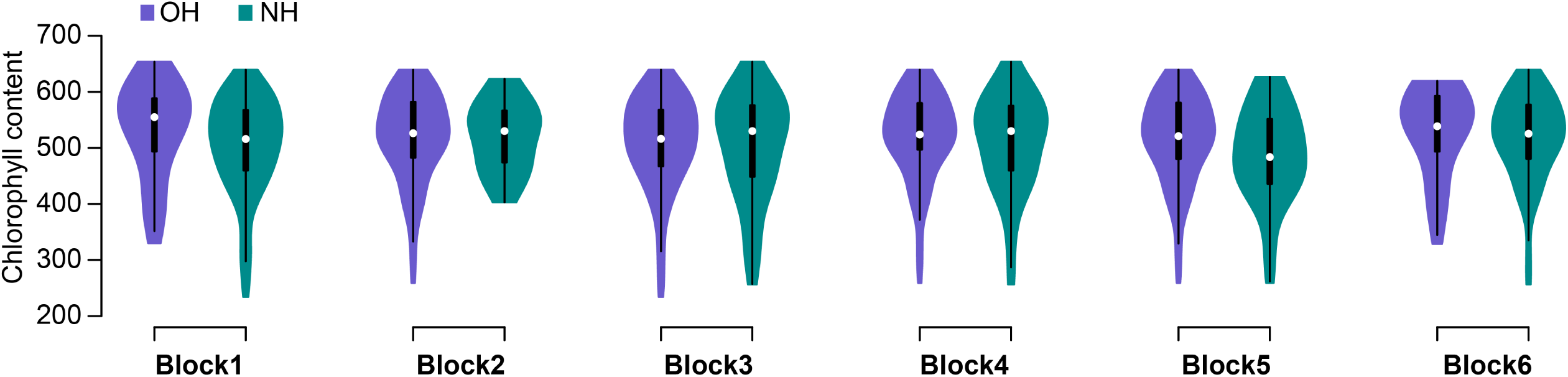
Phenotypic performance between Old-Era haplotype (OH) and New-Era haplotypes (NH) at Block1 to Block6 for leaf chlorophyll content.

**Figure S7.**
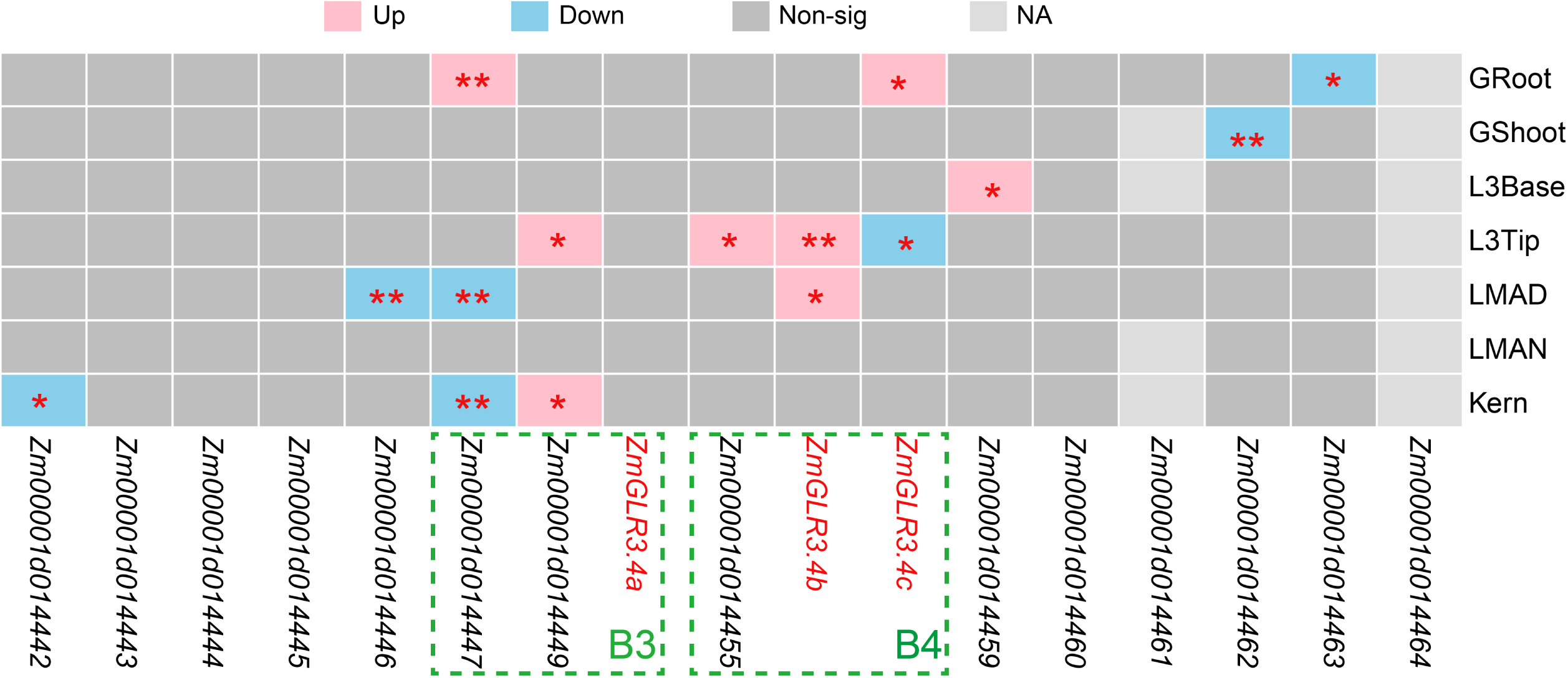
Gene expression differences between Old-Era and New-Era inbred lines for genes underneath the strongest selection signal at Chromosome 5. The pink and sky-blue colors indicate gene expression was upregulated and downregulated significantly in New-Era inbred lines, respectively. The dashed rectangles highlight genes located with LD block3 (B3) and block4 (B4). \**P*-value < 0.05 and \*\**P*-value < 0.01 from two-sided Student’s *t*-test.

**Figure S8.**
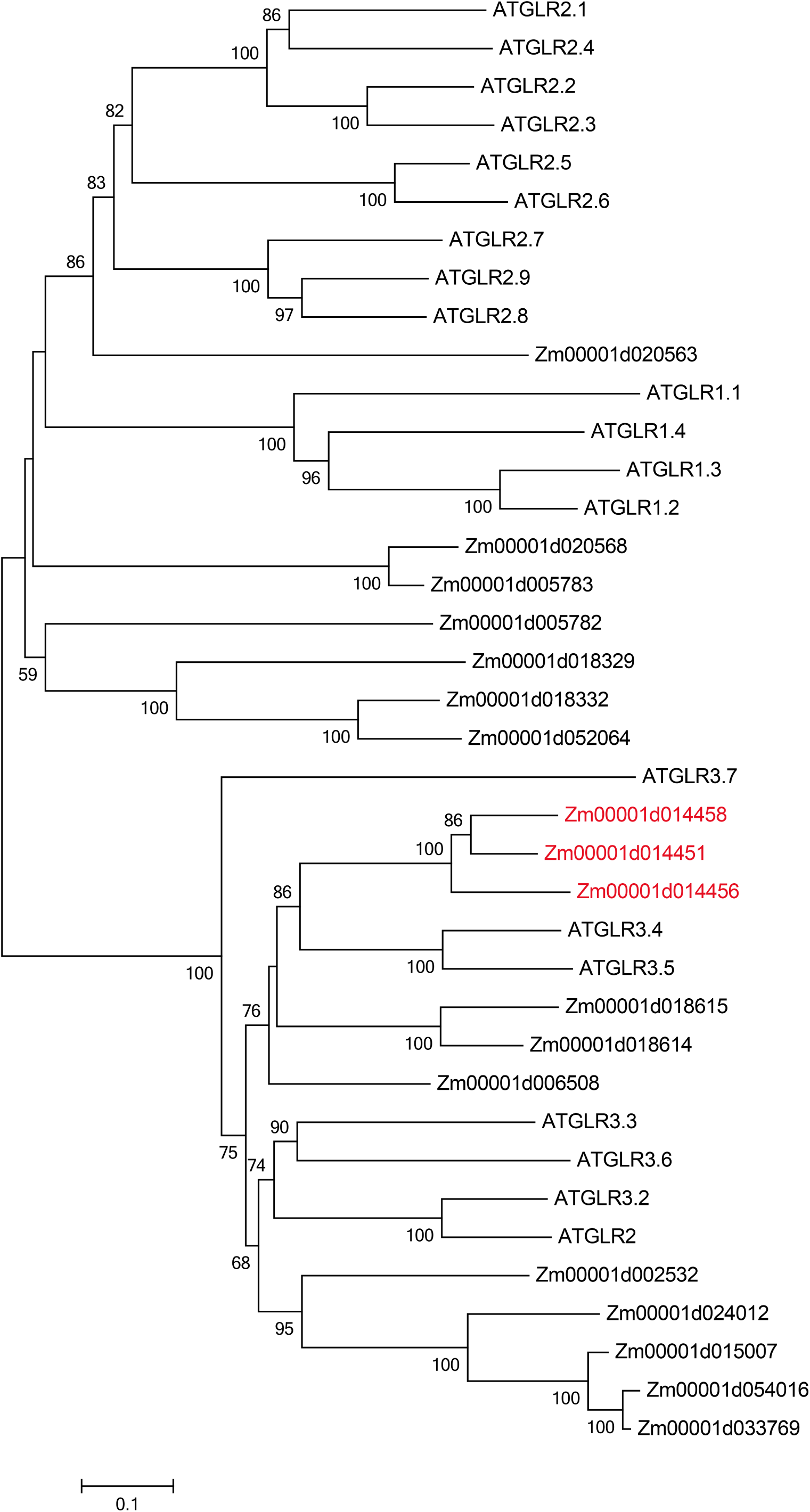
Phylogenetic tree generated from *Arabidopsis* and maize GLRs protein sequences. The gene names with red color indicate GLR genes under the strongest selection signal at chromosome 5.

**Figure S9.**
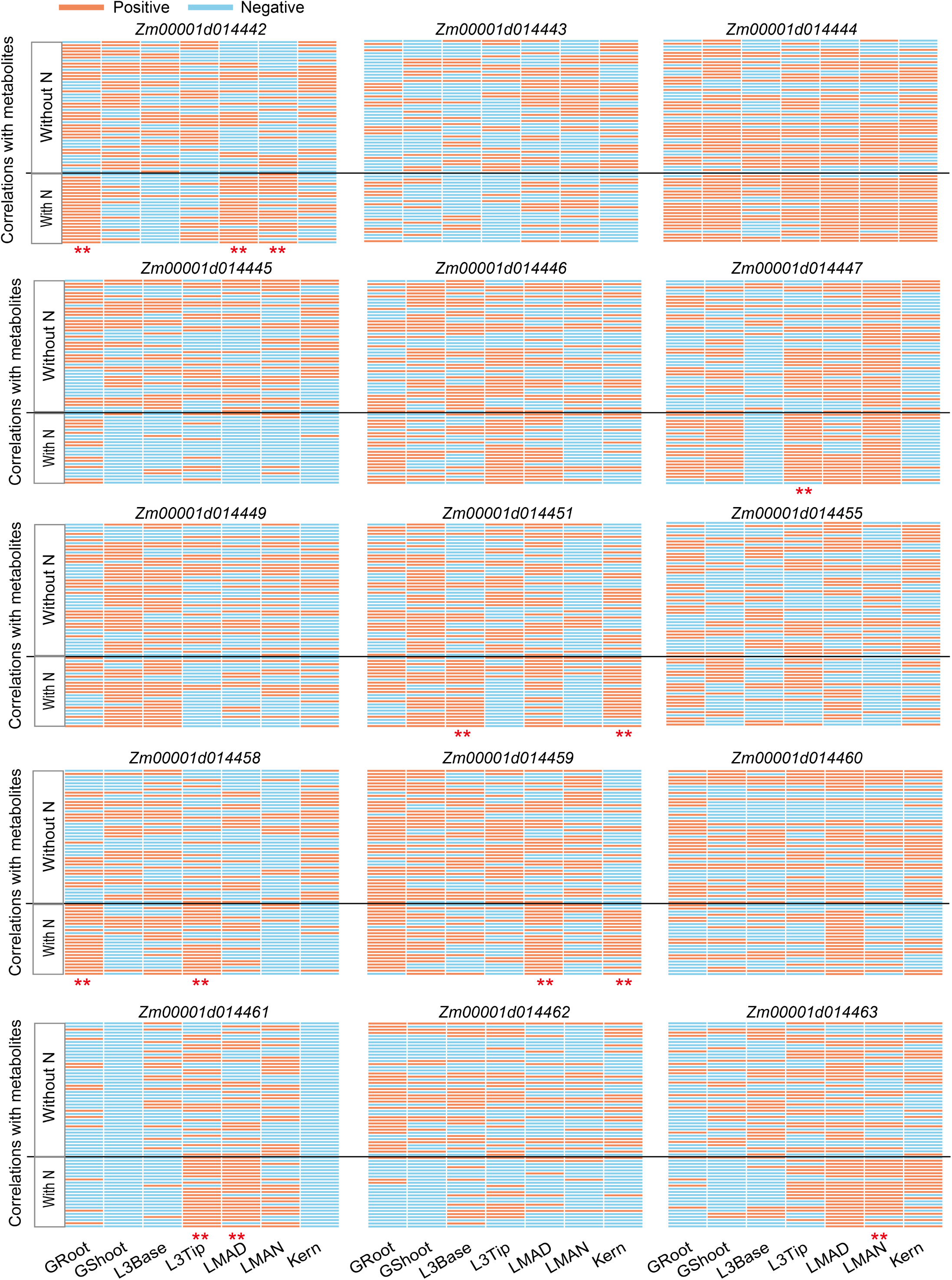
Correlation analysis between gene expression 66 metabolites. All these genes are located at the most significant sweep at Chromosome 5. “With N” and “Without N” denote the metabolites containing or not containing N in their formulas. ** denotes Chi-squared test *P*-value < 0.01.

**Figure S10.**
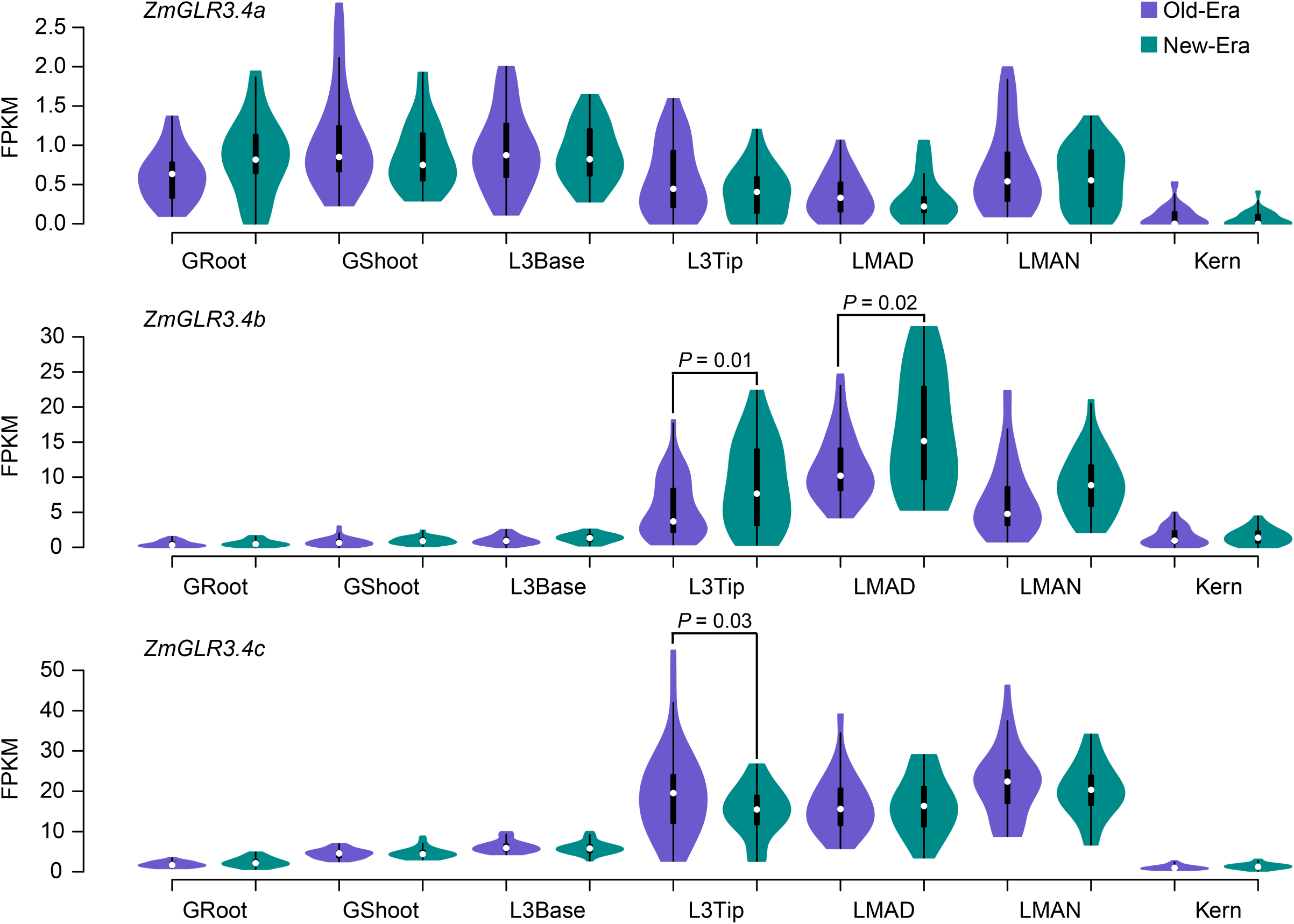
The distribution of gene expression levels of *ZmGLR3.4a, ZmGLR3.4b*, and *ZmGLR3.4c* in different tissues of Old- and New-Era inbred lines. *P* values were determined with a two-sided Student’s *t*-test. The data was obtained from Kremling et al. 2018.

**Figure S11.**
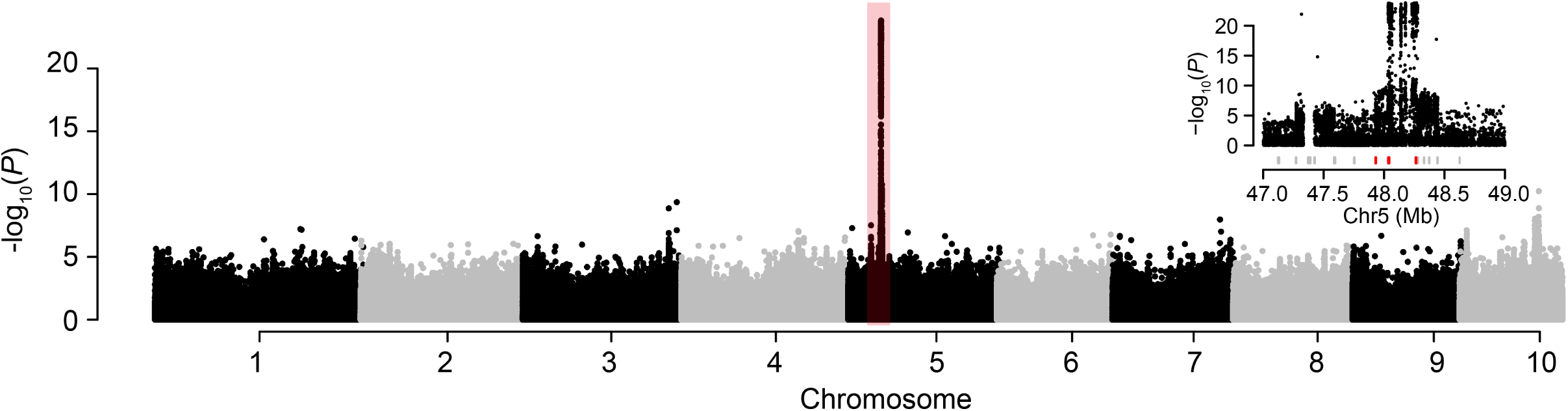
The Manhattan plot of the eQTL results using gene expression of *ZmGLR3.4c* collected from third leaf tip as the trait. The top right panel shows the zoom-in plot of the highlighted region in the Manhattan plot. The red ticks indicate the three GLR genes.

**Figure S12.**
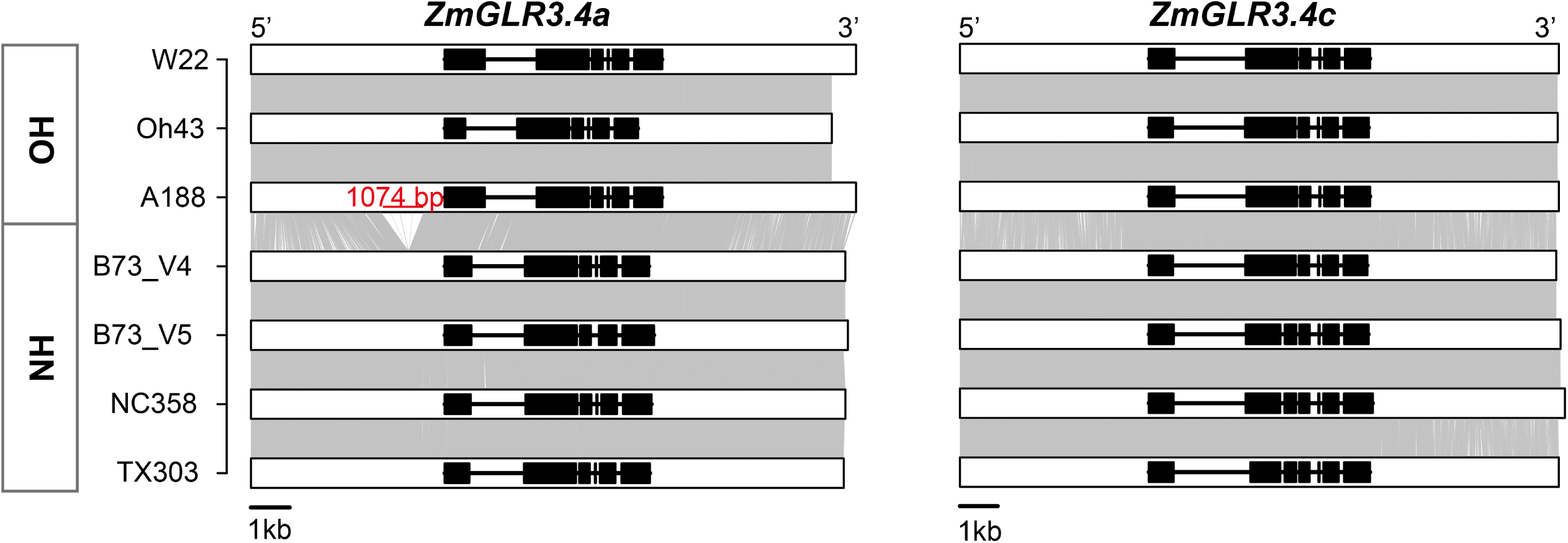
Comparison of *ZmGLR3.4a* and *ZmGLR3.4c* gene structures annotated from *de novo* assembled genomes. OH and NH indicate the inbred lines carrying the Old-Era and New-Era haplotype, respectively.

**Figure S13.**
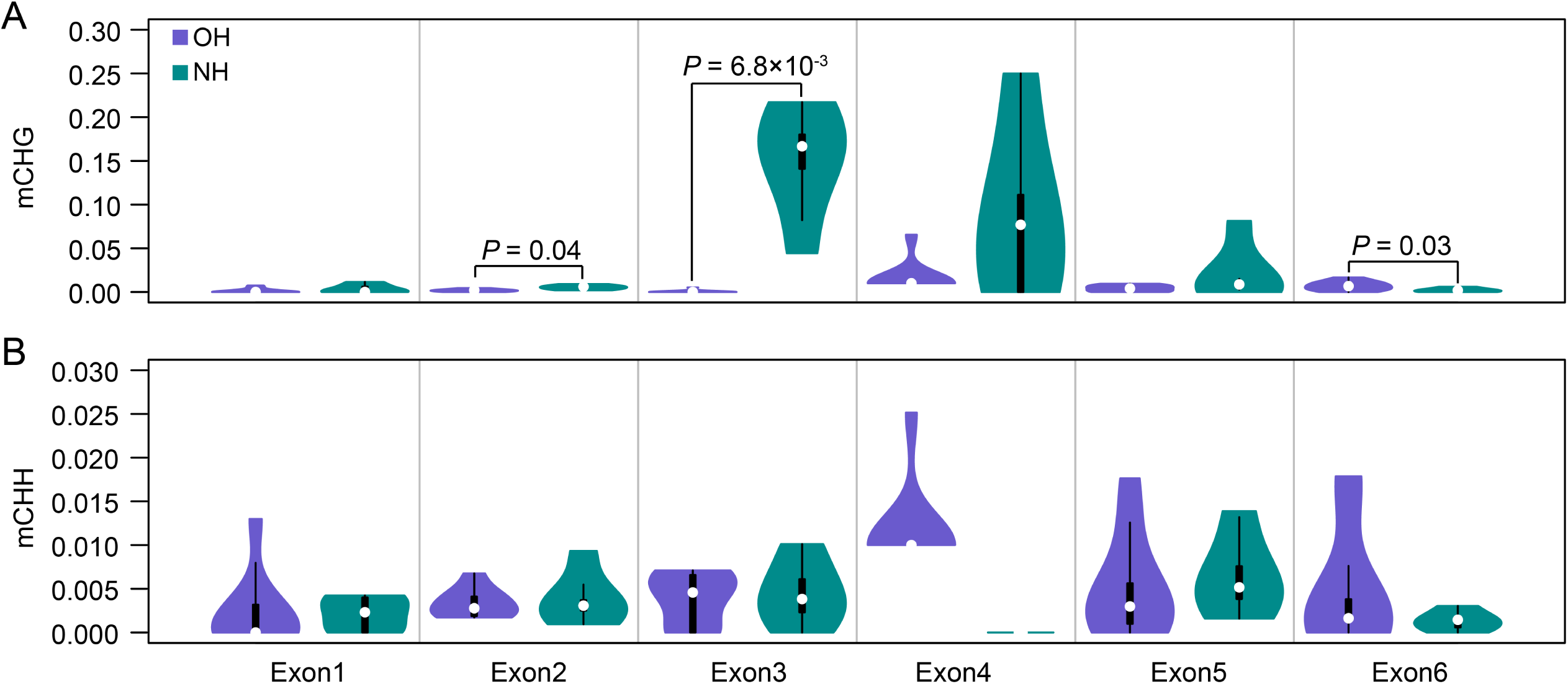
Levels of DNA CG (A) and CHG (B) methylation in exons of *ZmGLR3.4b*. OH and NH indicate the inbred lines carrying the Old-Era and New-Era haplotype, respectively. *P* values were determined with a two-sided Student’s *t*-test.

**Figure S14.**
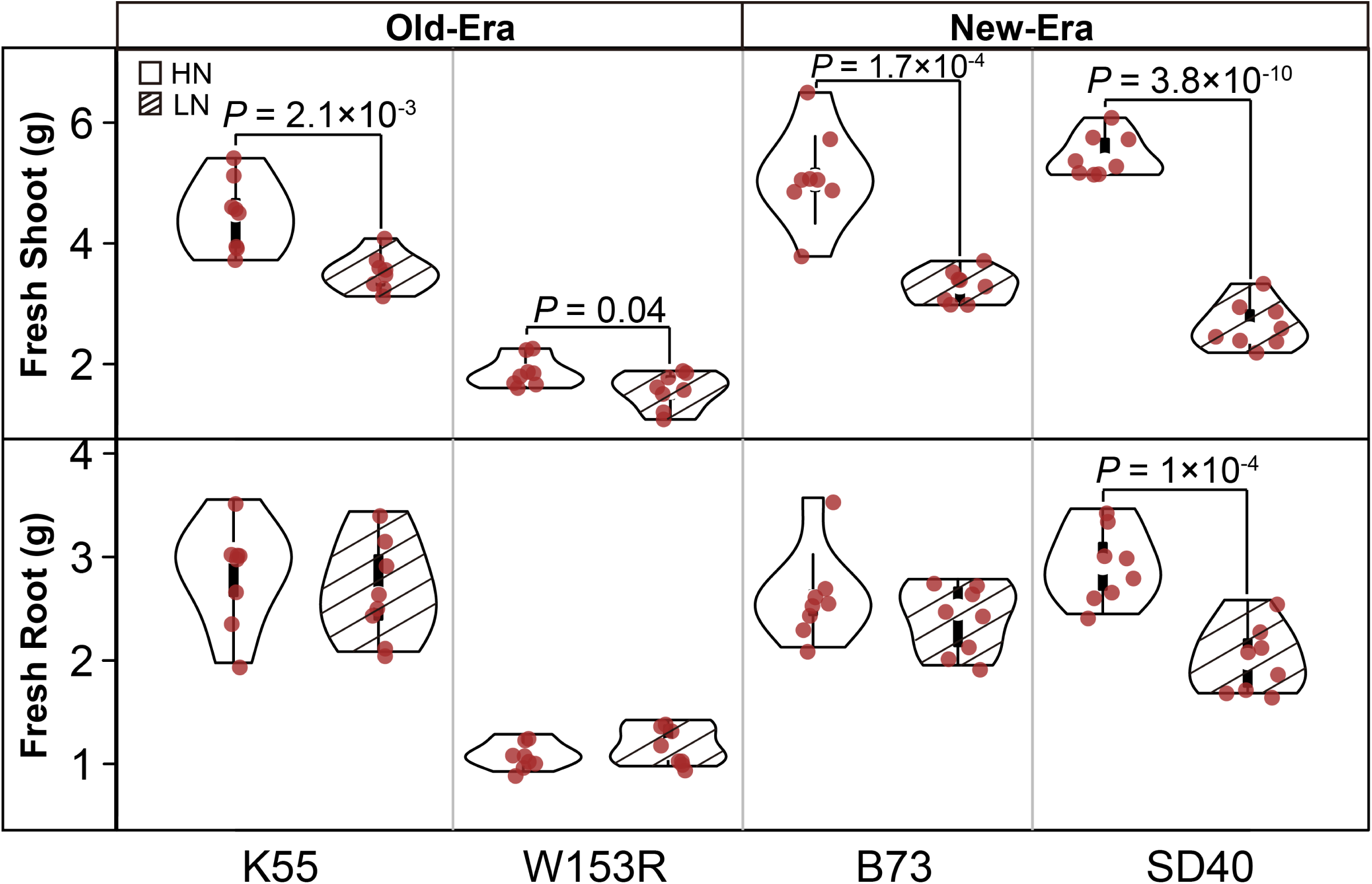
The fresh weight of shoot and root for Old-Era and New-Era inbred lines growing with high N and low N treatments. *P* values were determined with a two-sided Student’s *t*-test.

**Figure S15.**
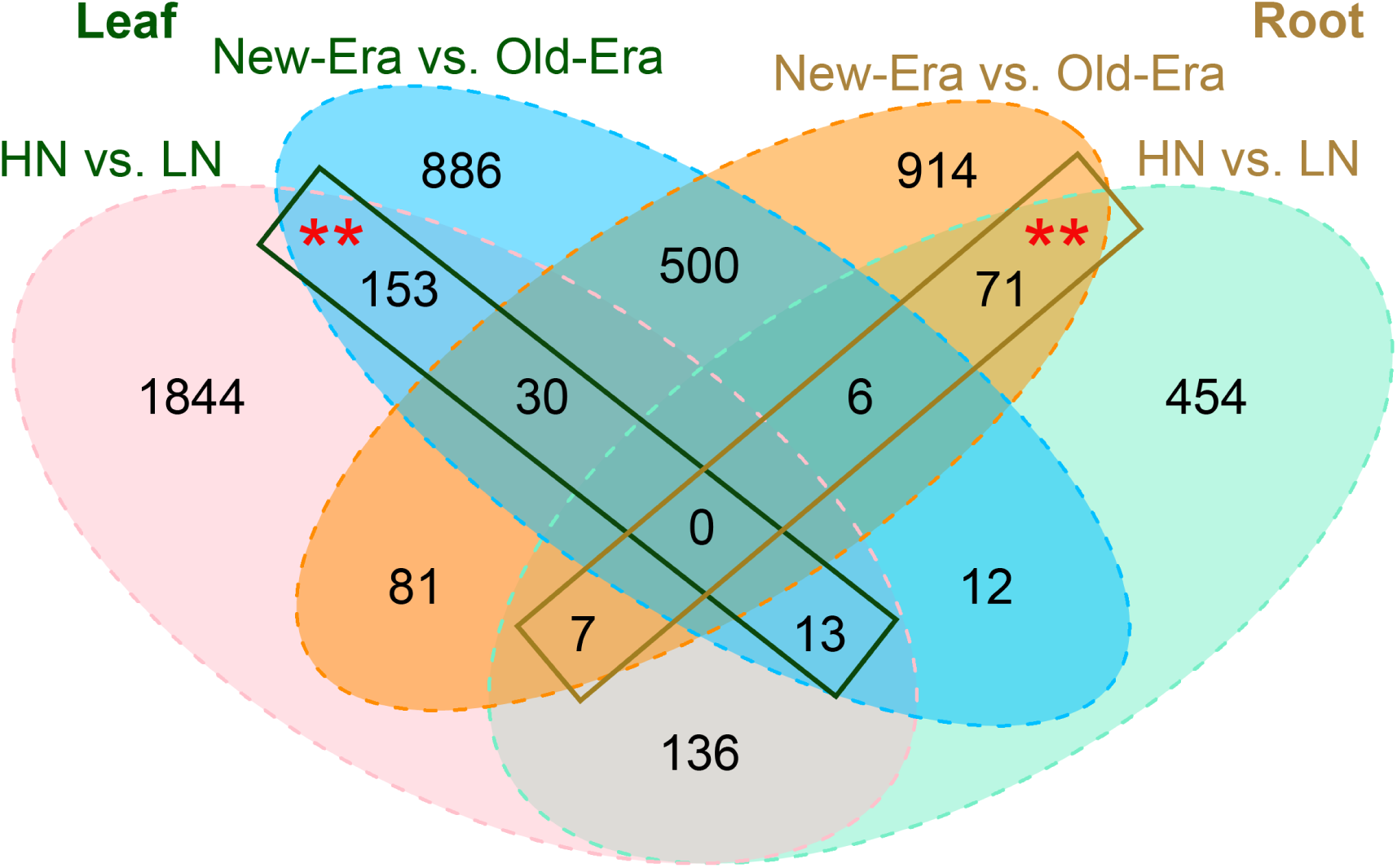
Differentially expressed (DE) genes between Old-Era and New-Era inbreds. Overlaps between the sets of DE genes of high N (HN) vs. low N (LN) and New-Era (NE) vs. Old-Era (OE) in leaf and root tissues. The red asterisks indicate categories with more genes than expected at a permutation based *P*-value cutoff of < 0.01.

**Figure S16.**
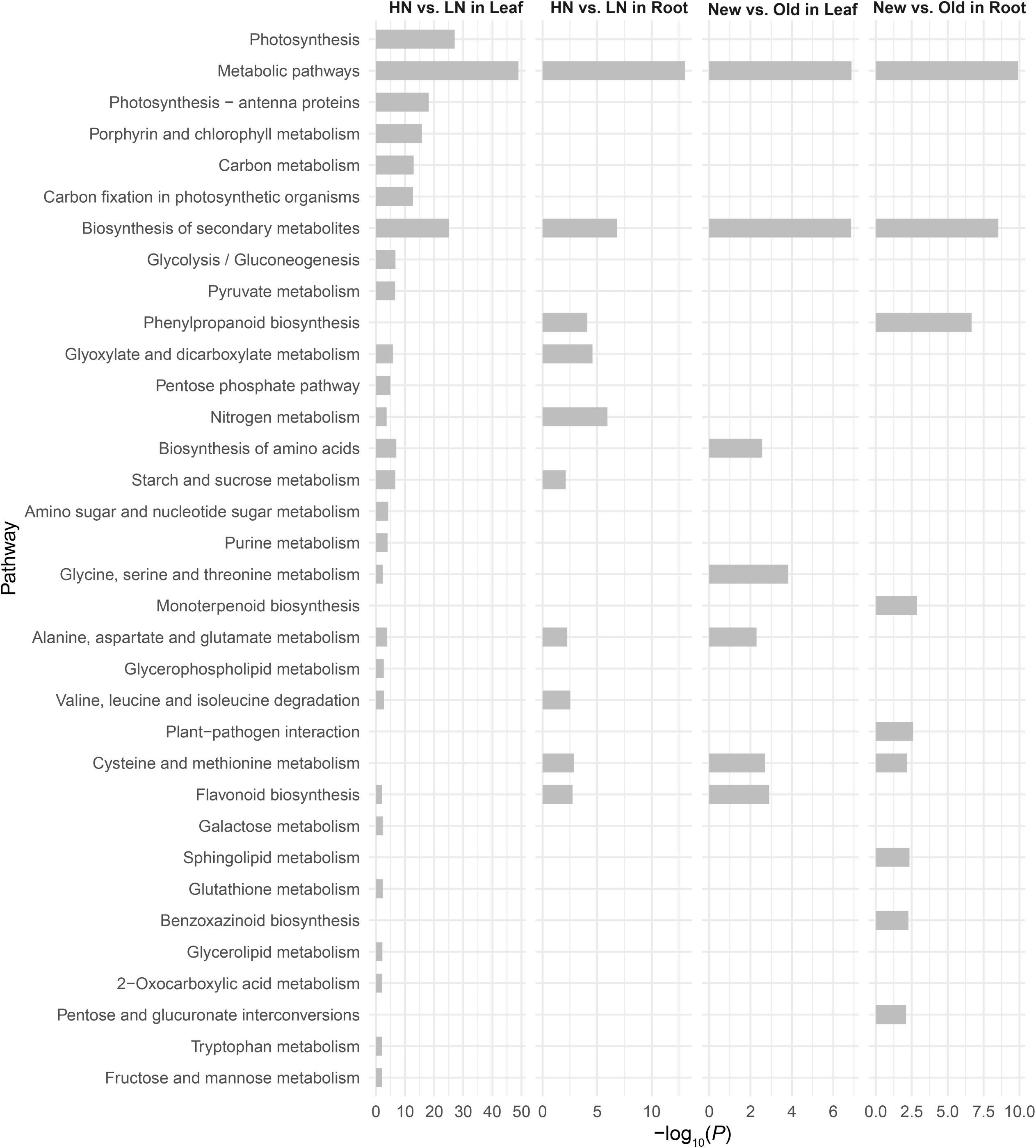
Enriched KEGG (Kyoto Encyclopedia of Genes and Genomes) pathways detected by N and Era responsive differentially expressed genes.

